# Cap-independent translation directs stress-induced differentiation of the protozoan parasite *Toxoplasma gondii*

**DOI:** 10.1101/2024.09.17.613578

**Authors:** Vishakha Dey, Michael J. Holmes, Matheus S. Bastos, Ronald C. Wek, William J. Sullivan

## Abstract

Translational control mechanisms modulate microbial latency of eukaryotic pathogens, enabling them to evade immunity and drug treatments. The protozoan parasite *Toxoplasma gondii* persists in hosts by differentiating from proliferative tachyzoites to latent bradyzoites, which are housed inside tissue cysts. Transcriptional changes facilitating bradyzoite conversion are mediated by a Myb domain transcription factor called BFD1, whose mRNA is present in tachyzoites but not translated into protein until stress is applied to induce differentiation. We addressed the mechanisms by which translational control drives BFD1 synthesis in response to stress-induced parasite differentiation. Using biochemical and molecular approaches, we show that the 5’-leader of *BFD1* mRNA is sufficient for preferential translation upon stress. The translational control of *BFD1* mRNA is maintained when ribosome assembly near its 5’-cap is impaired by insertion of a 5’-proximal stem-loop and upon knockdown of the *Toxoplasma* cap-binding protein, eIF4E1. Moreover, we show that a *trans*-acting RNA-binding protein called BFD2/ROCY1 is necessary for cap-independent translation of *BFD1* through its binding to the 5’-leader. Translation of *BFD2* mRNA is also suggested to be preferentially induced under stress, but by a cap-dependent mechanism. These results show that translational control and differentiation in *Toxoplasma* proceed through cap-independent mechanisms in addition to canonical cap-dependent translation. Our identification of cap-independent translation in protozoa underscores the antiquity of this mode of gene regulation in cellular evolution and its central role in stress-induced life-cycle events.

## Introduction

Cellular adaptation and differentiation processes in response to cellular stresses are initiated by a reprogramming of gene expression. A central feature of this reprograming is rapid lowering of global protein synthesis in favor of selective translation of a subset of mRNAs that include transcription factors that activate genes for stress remediation or differentiation (1–3). We have shown that this paradigm governs life cycle stage transitions in protozoan parasites that underlie transmission and pathogenesis of infectious disease (3–5). Study of translational control in these early-branching eukaryotes not only promises to reveal potential novel drug targets, but also sheds important mechanistic insights into the evolutionary origins of these regulatory processes (6–8).

*Toxoplasma gondii* is an obligate intracellular parasite of warm-blooded animals that causes opportunistic disease in humans. Feline species are the definitive host in which *Toxoplasma* undergoes its sexual stage, culminating in the dissemination of oocysts into the environment (9). Upon ingestion, the parasites transform into a rapidly replicative stage (tachyzoite) that disseminates throughout the host body before converting into a quiescent stage (bradyzoite) housed in tissue cysts that can be sustained for the life of the host. Tissue cysts also provide another major route of transmission through the consumption of raw or undercooked meat (10). Reactivation of bradyzoites can occur when patients become immunocompromised, producing life-threatening damage to critical areas where tissue cysts typically reside, such as the brain, heart, and skeletal muscle (11). Current treatments of toxoplasmosis only target replicating tachyzoites and do not appear to exert appreciable activity against the formation or viability of tissue cysts (12). The formation of latent tissue cysts allows *Toxoplasma* to persist in its host and prevents a radical cure of this infection.

Given the importance of bradyzoites in driving parasite transmission and chronic disease, much research is aimed at understanding the molecular mechanisms orchestrating the conversion between tachyzoites and bradyzoites (13). Differentiation between life cycle stages requires the reprogramming of gene expression, while chromatin remodeling machinery and several apetala-2 (AP2) factors have been shown to contribute to the transcription of bradyzoite-inducing genes (14). Recently, Lourido and colleagues identified a “master regulator” transcription factor termed BFD1 (Bradyzoite Formation Deficient-1) that is necessary and sufficient for bradyzoite formation (15).

BFD1 is a SANT/Myb-like DNA-binding protein whose mRNA is present in tachyzoites but not translated until parasites are exposed to a bradyzoite-induction stress (15). We previously suggested with polysome profiling that *BFD1* mRNA is preferentially translated when tachyzoites are subjected to a stress that induces bradyzoite differentiation (16). These studies indicate that expression of BFD1 protein is predominantly regulated by translational control and the elevated levels of BFD1 are sufficient to confer transcription events directing cyst formation. Consistent with this idea, the 5’-leader sequence of *BFD1* mRNA is about 2.7 kb in length, harboring multiple predicted upstream open reading frames (uORFs), which are known in other model systems to regulate start codon selection and translation efficiency of the protein coding sequence (CDS) (17). Moreover, a CCCH-type zinc finger mRNA-binding protein called BFD2 (also referred to as ROCY1) has recently been described that is suggested to direct the stress-dependent translation of *BFD1* (18, 19).

Bulk cellular translation begins with eIF4F association with the 5’-cap of mRNAs through its eIF4E subunit, which then recruits the ribosomal preinitiation complex that proceeds to scan the 5’-leader for an optimal start codon (20). This cap-dependent translation can be modulated by uORFs that can be bypassed by scanning ribosomes or allow for re-initiation for efficient CDS translation (21). Cap-independent mechanisms have also been described, most commonly among viruses and less frequently for cellular mRNAs (22–24). Translation that can occur independent of eIF4E cap-association and the subsequent ribosome scanning typically involves direct entry of ribosomes onto the 5’-leader sequence by secondary structures that interface with *trans*-acting proteins to recruit initiating ribosomes (23, 24).

In this study, we addressed the mechanisms of preferential translation of *BFD1* during stress-induced bradyzoite differentiation. Utilizing luciferase reporter assays and genetically modified *Toxoplasma* parasites, we show that *BFD1* translation can occur through a cap-independent mechanism that involves BFD2 binding to the 5’-leader sequence. By comparison, preferential translation of *BFD2* during stress occurs by cap-dependent processes. Our results represent the first evidence of cap-independent translation in protozoa, suggesting that it is an ancient mechanism of gene regulation present in early-branching eukaryotes and is critical for directing differentiation in the *Toxoplasma* life-cycle.

## Results

### 5’-leader sequence of *BFD1* mRNA regulates its translation

It has been reported that *BFD1* mRNA levels are similar in tachyzoites and developing bradyzoites (15); moreover, *BFD1* knockout parasites containing an ectopic copy of *BFD1* only express detectable BFD1 during bradyzoite differentiation. These earlier findings suggested that *BFD1* mRNA is preferentially translated during stress conditions. Supporting this premise, our earlier polysome profiling study suggested that *BFD1* transcripts are shifted significantly towards translating ribosomes during bradyzoite-induction stress (16). To further support stress-induced preferential translation of *BFD1*, we endogenously tagged the genomic locus to fuse an HA epitope onto the C-terminus of BFD1 protein in cystogenic wild-type (WT) ME49 strain parasites (BFD1^HA^, Fig. S1A). Immunoblotting revealed expression of the tagged BFD1^HA^ protein with the predicted size (262 kDa) only after parasites were exposed to alkaline stress, a commonly used bradyzoite induction trigger (Fig. S1B). An immunofluorescence assay (IFA) verified stress-dependent BFD1^HA^ protein expression in the parasite nucleus (Fig. S1C). Proper tagging of the *BFD1* gene was further confirmed by using specific primers and PCR analysis of genomic DNA (Fig. S1D).

To determine the stress-dependent expression patterns for *BFD1* mRNA and encoded protein, we conducted a time course of alkaline stress exposure in BFD1^HA^ parasites. *BFD1* mRNA was detectable in unstressed tachyzoites, and transcript levels remained unchanged in response to 24 h alkaline stress (Fig. 1A). Parasites subjected to prolonged alkaline stress (5 days) displayed a modest but statistically significant increase in *BFD1* mRNA, which is consistent with a report suggesting that *BFD1* regulates its own transcription after 48 h alkaline stress (15). As a control, we showed that a well-documented marker of bradyzoite differentiation, *BAG1*, was increased more than 50-fold after 24 h of alkaline stress, with further enhancement after 5 days (Fig. 1A). By comparison, SAG1, known to be expressed only in tachyzoites, was potently lowered at 24 h of stress, with minimal transcript after 5 days. Despite the presence of *BFD1* mRNA, BFD1^HA^ protein was not detectable until tachyzoites were treated with the bradyzoite inducing stress for 24 h, with further increase after 5 days (Fig. 1B). Consistent with the observed robust enhancement of *BAG1* mRNA, BAG1 protein was also only well-expressed in the stressed parasites (Fig. 1B). These results support the model that *BFD1* mRNA is preferentially translated in response to a stress that elicits *Toxoplasma* differentiation.

**Figure 1.**
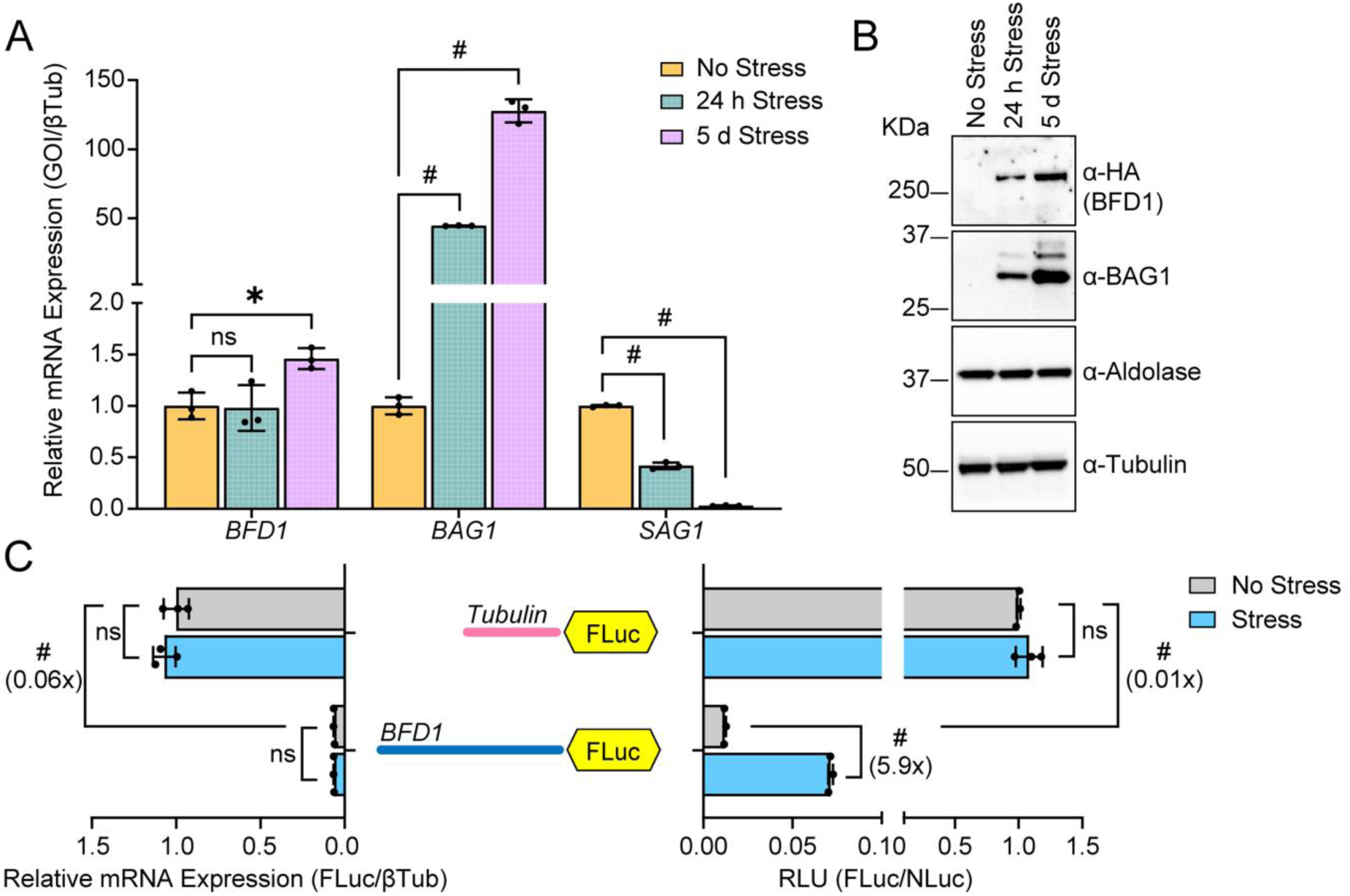
*BFD1* mRNA is preferentially translated under alkaline stress. BFD1^HA^ parasites were cultured in tachyzoite conditions (No Stress) or under alkaline stress conditions (Stress) for 24 h or 5 days. (A) cDNA was synthesized from total RNA collected from the parasites and mean transcript abundance ± standard deviation from 3 biological replicates is plotted relative to constitutive expressed β-Tubulin. Values are normalized to no stress tachyzoites for each transcript. ns = p>0.05; * = p≤0.01, # = p≤0.0001 by Student’s two-tailed t-test. (B) Immunoblot of lysates from unstressed and stressed BFD1^HA^ parasites were probed with the designated antibody. Molecular weight markers are indicated to the left of each panel in kDa. (C) Tub-FLuc or BFD1-FLuc reporters are highlighted in the illustration including the 5’-leaders of Tubulin and BFD1 (not to scale) fused to firefly luciferase (FLuc). Reporters were co-transfected with Tub-NLuc into WT ME49 parasites and cultured under normal no stress or alkaline stress conditions for 24 h. Luciferase measurements were obtained by Dual Glo luciferase assay in alkaline stress (Stress, blue bars) or tachyzoite conditions (No Stress, grey bars). All luciferase measurements were adjusted into relative luciferase units (RLUs) by normalizing against Tub-FLuc reporter in unstressed culture conditions ± standard deviation from 3 biological replicates, which is plotted on right. Luciferase reporter mRNAs shown on the left of the panels were measured by RT-qPCR. Mean transcript abundance ± standard deviation from 3 biological replicates is plotted relative to β-Tubulin, with normalization to tachyzoites transfected with Tub-FLuc reporter. ns = p>0.05; # = p≤0.0001 by Student’s two-tailed t-test. Fold changes are shown in parentheses.

To determine whether the unusually long 5’-leader of the *BFD1* mRNA is sufficient for driving preferential translation, we positioned it upstream of a firefly luciferase (FLuc) reporter driven by the constitutive tubulin promoter. The first 35 codons of the *BFD1* CDS were also included to maintain an out-of-frame uORF. This reporter, designated BFD1-FLuc, was transiently transfected into WT ME49 parasites in combination with a plasmid encoding constitutively expressed Nano luciferase (NLuc) to serve as a normalization control. A dual luciferase activity assay was performed on unstressed tachyzoites and tachyzoites subjected to 24 h alkaline stress to induce bradyzoite differentiation. We verified that both NLuc activity and its mRNA expression did not appreciably change in response to alkaline stress (Fig. S2A-B). Using this reporter system, we found that alkaline stress induces a ~6-fold increase in BFD1-FLuc activity, with no significant difference in the reporter mRNA as quantified by RT-qPCR (Fig. 1C and S2A). By comparison, a control reporter consisting of the 5’-leader sequence for tubulin (Tub-FLuc) displayed no change in activity between unstressed and stressed conditions (Fig. 1C and S2A-B). These results indicate that the *BFD1* 5’-leader sequence is sufficient to direct stress-induced preferential translation.

### Stress-dependent translation of *BFD1* mRNA is regulated by a cap-independent mechanism

*BFD1* is transcribed at a single transcriptional start site (25, 26), generating an unusually long 5’-leader that is replete with predicted uORFs (Fig. S3A). In other model systems, uORFs have been shown to play a major role in canonical cap-dependent translation by usurping scanning ribosomes and regulating translation at the downstream CDS (1, 17, 27). To determine whether cap-binding and ribosomal scanning of the 5’-leader is required for the preferential translation of *BFD1* during stress, we inserted a stable stem-loop structure just downstream of the 5’-end of the reporter mRNA constructs. Stable secondary structures, such as this stem-loop, in proximity of the 5’-cap lower the ability of ribosomes to bind the 5’-ends of mRNAs and initiate scanning for initiation codons (28). As expected, insertion of the stem-loop into Tub-FLuc largely abrogated luciferase activity, indicating that translation of Tub-FLuc is cap-dependent (Fig. 2A). By contrast, while insertion of the stem-loop into BFD1-FLuc lowered basal translation, it did not diminish the ~6-fold increase in luciferase activity in response to stress (Fig. 2B). Comparable mRNA levels of the luciferase reporters were observed independent of stress, indicating that changes in BFD1-Fluc activity were the result of translational control. These findings suggest that induced translation of *BFD1* mRNA in response to stress occurs by a mechanism that is not dependent on the 5’-cap.

**Figure 2.**
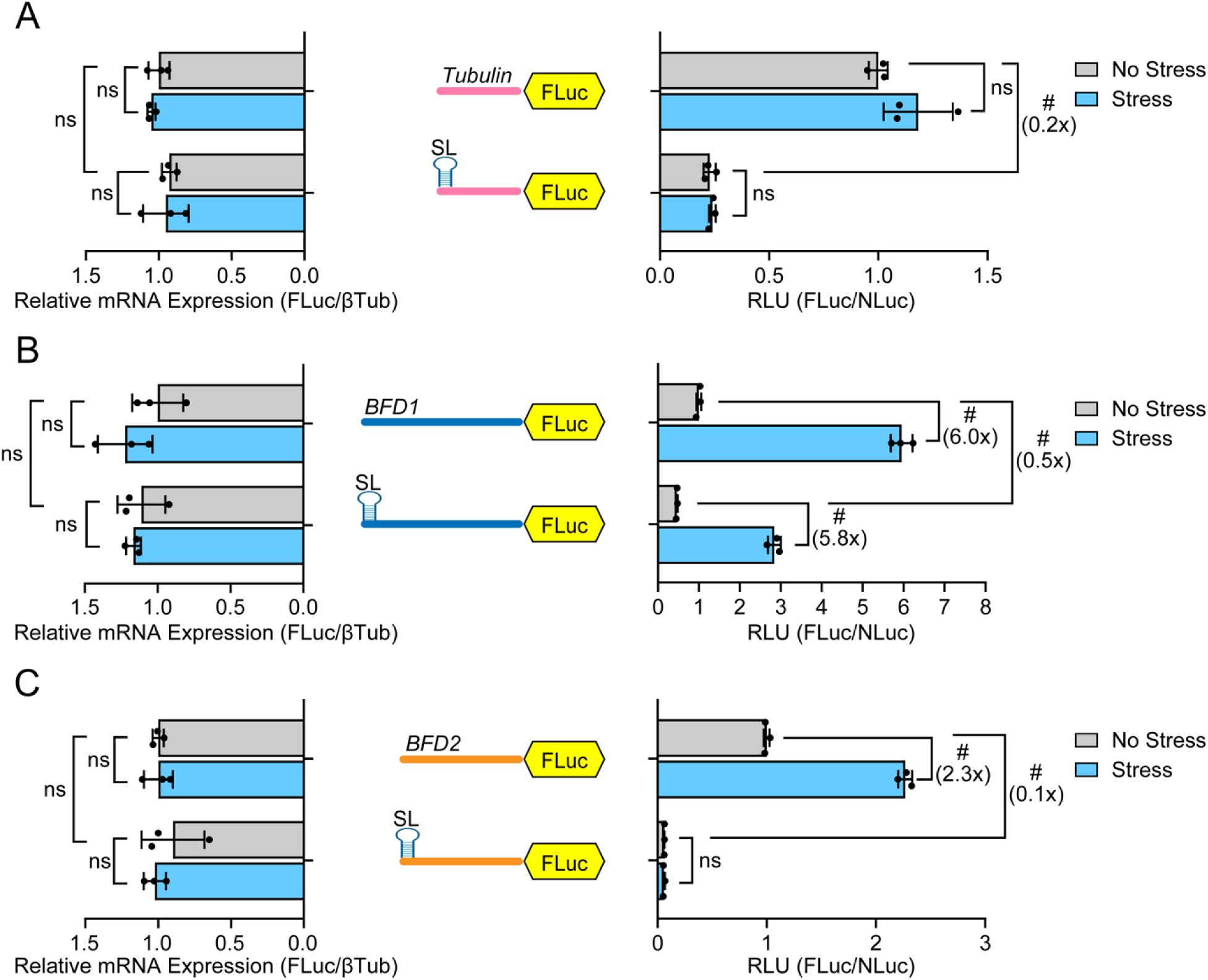
Insertion of stem-loop blocks preferential translation of *BFD2* but not *BFD1*. The Tub-FLuc (A), BFD1-FLuc (B) and BFD2-Fluc (C) reporters featuring the respective 5’-leaders adjoined to the FLuc coding sequence with or without the indicated 5’-stem-loop (SL). These reporters were individually co-transfected with Tub-NLuc into WT ME49 parasites. RLU was measured for transfected parasites cultured under alkaline stress (Stress, blue bars) or tachyzoite conditions (No Stress, grey bars). Mean fold change ± standard deviation from 3 biological replicates is plotted with normalization to the RLU of transfected parasites without the stem-loop with no stress. Luciferase reporter mRNAs measured by RT-qPCR. Mean transcript abundance ± standard deviation from 3 biological replicates is plotted relative transfected tachyzoites without the stem-loop with no stress. ns = p>0.05; # = p≤0.0001 by Student’s two-tailed t-test. Fold changes are shown in parentheses.

To examine whether cap-independent mechanisms operate on other developmentally related genes in *Toxoplasma*, we investigated the regulation of BFD2/ROCY1, a CCCH zinc-finger protein that promotes *BFD1* expression by binding its mRNA in a stress-dependent manner (18, 19). Like BFD1, expression of BFD2 protein is enhanced during stress-induced bradyzoite induction (18). We generated a BFD2 luciferase reporter comprised of the 5’-leader of *BFD2*, which is ~750 nt in length, expressed by the tubulin gene promoter (BFD2-FLuc). The length of the 5’-leader we tested was chosen based on a reported major eIF4E1 binding site most proximal to the CDS as determined by CLIP-seq (Fig. S3B) (26). Similar to BFD1-FLuc, 24 h alkaline stress caused a ~2.3-fold increase in BFD2 luciferase activity, with minimal changes in mRNA levels (Fig. 2C). However, distinct from BFD1-FLuc, the basal expression and stress-dependent upregulation were largely abolished when the stem-loop was present (Fig. 2C). These results suggest that *BFD2* mRNA can be preferentially translated in response to stress but by a cap-dependent mechanism.

### Depletion of eIF4E1 does not inhibit preferential translation of *BFD1* in response to stress

To further evaluate cap-dependency for *BFD1* translational control, we assessed the consequence of eIF4E1 depletion on BFD1-FLuc activity induced by stress. We reported that *Toxoplasma* eIF4E1 is well-expressed in tachyzoites, associates with the 5’-end of mRNAs genome-wide, and is critical for global translation (26). In this earlier study, we generated a conditional knockdown parasite line in ME49 strain that fused eIF4E1 to a minimal auxin-inducible degron (mAID) tag, designated eIF4E1^mAID-HA^ (26). These eIF4E1^mAID-HA^ parasites contained an integrated FLuc gene that we knocked out by inserting an HXGPRT selectable marker cassette, thereby generating eIF4E1^mAID-HAΔFLuc^ parasites that could be used with our reporter constructs. Assays confirmed the loss of luciferase activity in eIF4E1^mAID-HAΔFLuc^ parasites (Fig. S4A). The auxin-mediated depletion of eIF4E1 as judged by immunoblot was unaffected by removal of the integrated FLuc gene, with near complete loss of eIF4E1 after 4 h treatment with 500 μM auxin 3-indoleacetic acid (IAA) (Fig. S4B).

We next compared the luciferase activity of our reporter constructs under stressed and unstressed conditions in eIF4E1^mAID-HAΔFLuc^ parasites followed by 4 h treatment with IAA or vehicle (DMSO) control. We noted that NLuc itself is sensitive to eIF4E1 depletion, confounding the ability to normalize samples; therefore, raw FLuc data and accompanying mRNA measurements are presented for this reporter series (Fig. S5). Once again, we detected stress-induced translational expression of BFD1-FLuc and BFD2-FLuc, but not for the Tub-FLuc control (Fig. 3A-C). Notably, loss of eIF4E1 abrogated the stress-dependent increase in translation of BFD2-FLuc but not BFD1-FLuc (Fig. 3B-C). As noted earlier with the BFD1 reporter featuring the 5’-positioned stem-loop (Fig. 2B), there was some reduction in basal expression upon eIF4E1 depletion by IAA, but the >5-fold induction by stress was retained (Fig. 3B). Furthermore, the combined insertion of the 5’-stem-loop and IAA treatment showed a similar robust induction (Fig. 3B). There were minimal changes in mRNA levels for the respective reporters in the presence or absence of IAA-directed depletion of eIF4E1, indicating that changes in luciferase activities between stressed and unstressed samples were not due to differences in relative mRNA expression (Fig. 3A-C). These results further support the model that preferential translation of *BFD1* during stress occurs by a mechanism not dependent on the 5’-cap, whereas *BFD2* employs a canonical cap-dependent mechanism.

**Figure 3.**
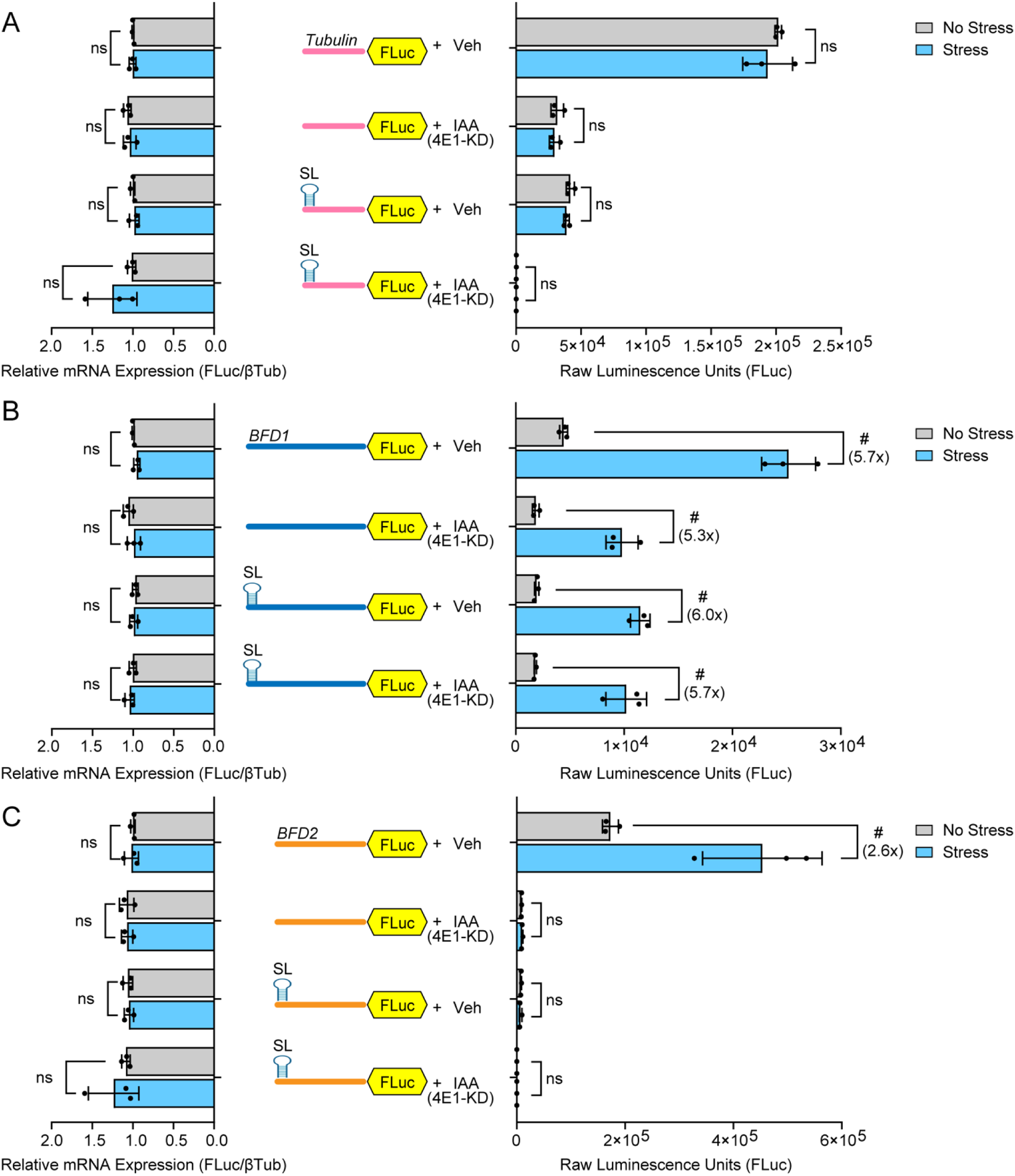
Translation of *BFD1* persists despite depletion of eIF4E1. (A), Tub-FLuc reporter with or without the stem-loop co-transfected with Tub-NLuc into eIF4E1^mAID-HAΔFLuc^ parasites. Transfected parasites were cultured under alkaline stress (Stress, blue bars) or tachyzoite conditions (No Stress, grey bars) for 24 h. Parasites were treated with vehicle (Veh) or IAA for 4 h for targeted depletion of eIF4E1. FLuc activity was measured and mean raw luminescence units ± standard deviation from 3 biological replicates is presented normalized to Tub-FLuc without stress (Vehicle). ns = p>0.05; # = p≤0.0001 by Student’s two-tailed t-test. Luciferase reporter mRNAs were also measured by RT-qPCR as described in Fig 2. ns = p>0.05 by Student’s two-tailed t-test. Fold changes relative to unstressed parasites are shown in parentheses. BFD1-FLuc (B) and BFD2-FLuc (C) reporters with or without the stem-loop co-transfected with Tub-NLuc into eIF4E1^mAID-HAΔFLuc^ parasites and luciferase activity and mRNA measurements and results are presented as in panel A.

### BFD2-binding sites on *BFD1* mRNA facilitate preferential translation

We previously reported genome-wide CLIP-seq of endogenously tagged BFD2, which revealed three BFD2-binding sites on the BFD1 mRNA 5’-leader that we designated BR1, BR2, and BR3 (for BFD2 <u>B</u>inding <u>R</u>egion) (Fig. 4A) (19). We hypothesized that BFD2 may contribute to the cap-independent mechanism directing preferential translation of *BFD1*. To test this idea, we generated a series of BFD1-FLuc reporters containing individual deletions or combinations of these BFD2-binding sites. Deletion of all three BFD2-binding regions together in the 5’-leader of the BFD1-FLuc construct largely abrogated the stress-induced translation of BFD1-FLuc (Fig. 4B). While deletion of BR1 alone lowered basal expression of the reporter, it retained the ~6-fold increase in BFD1-FLuc activity following alkaline stress (Fig. 4C). In contrast, deletion of BR2 or BR3 alone partially lowered the stress-induced BFD1-FLuc to only 3-fold (Fig. 4C). There were no appreciable changes in mRNA levels for these reporters independent of stress, supporting that the measured stress-induction resulted from translational control. These results support the model that BFD2 binding to the 5’-leader of *BFD1* mRNA plays a critical role in its preferential translation during stress, with BR2 and BR3 being the predominant contributors.

**Figure 4.**
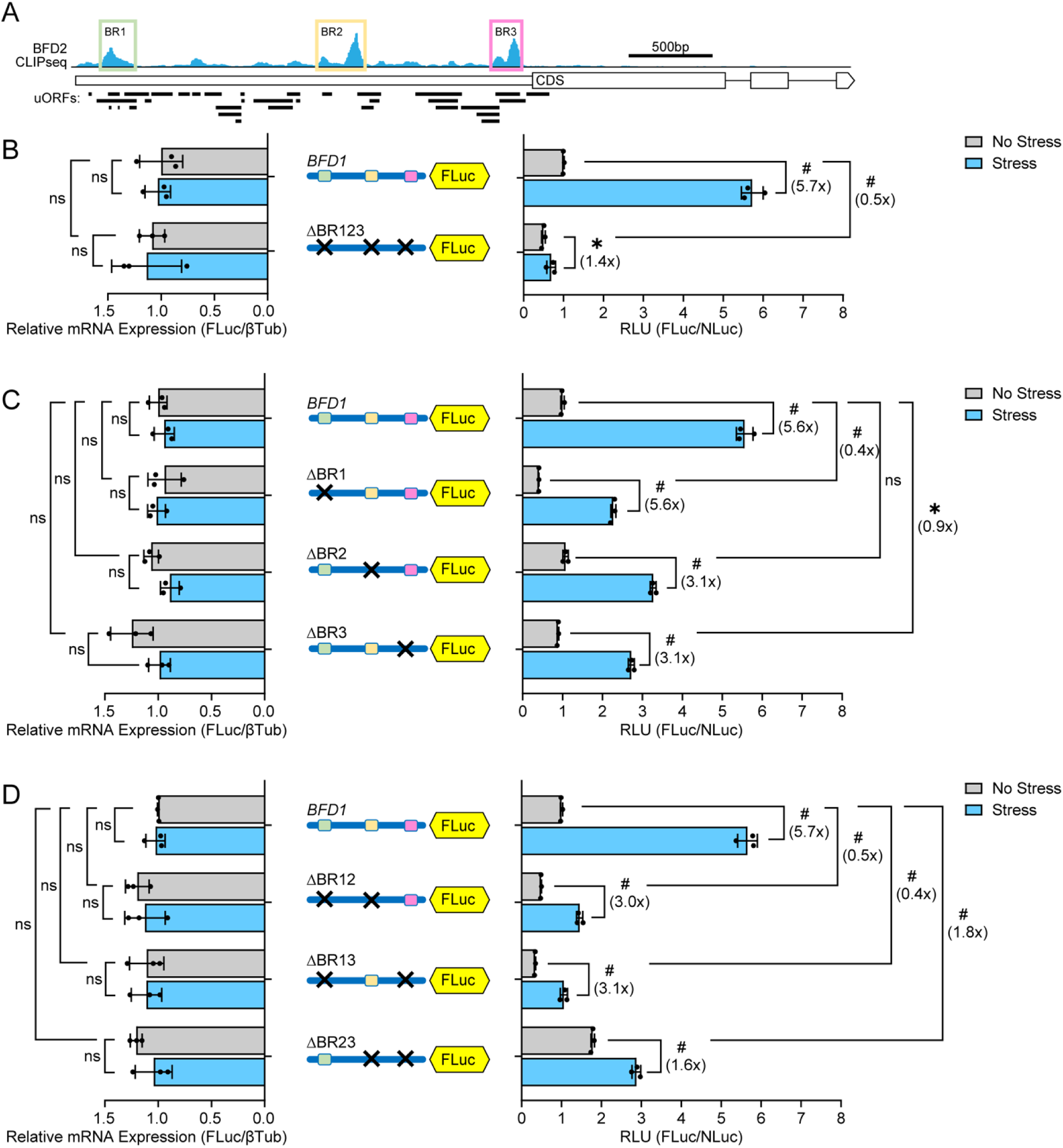
Preferential translation of *BFD1* requires BFD2-binding elements in its 5’ leader. (A) Schematic of BFD2 CLIP-seq highlighting BFD2-binding regions within the 5’-leader of the *BFD1* transcript. Illustrations in subsequent panels depict each BFD2-binding site as a different colored box with “X” denoting the deletion of that BFD2-binding site. (B) Unmodified and mutant *BFD1* 5’-leader versions with all BFD2-binding sites deleted were transfected into WT ME49 parasites. Transfected parasites were cultured under alkaline stress (Stress, blue bars) or tachyzoite culture conditions (No Stress, grey bars) for 24 h and relative luciferase activity was measured. Mean fold change ± standard deviation from 3 biological replicates is plotted with normalization to the RLU of the WT BFD1-FLuc reporter with no stress. Luciferase reporter mRNAs were measured by RT-qPCR. Mean transcript abundance ± standard deviation from 3 biological replicates is plotted relative to β-Tubulin with normalization to no stress transfected with BFD1-FLuc reporter. ns = p>0.05; * = p≤0.01; # = p≤0.0001 by Student’s two-tailed t-test. Fold changes are shown in parentheses. (C-D) Intact BFD1-FLuc reporter or with individual deletions (C) of BFD2-binding sites or combinations of these deletions (D) were co-transfected with Tub-NLuc into WT ME49 parasites. Luciferase activity and mRNA measurements and results are presented as in panel B.

We next tested the effects of pairwise deletions of the BFD2-binding sites for the *BFD1* translational control. Deletion of BR1+BR2 or BR1+BR3 each resulted in a 3-fold induction of BFD1-FLuc activity during stress (Fig. 4D), which represented no change from the individual deletions for BR2 and BR3, respectively. In contrast, deletion of BR2+BR3 nearly ablated stress-induced translation of BFD1-FLuc (Fig. 4D). The importance of the BR2 and BR3 sites for BFD1 translational control was also reinforced with individual and combination deletions of the BFD2 binding sites in a version of the BFD1-FLuc reporter with the inhibitory 5’-stem-loop as described earlier (Fig 5). In the BFD1-FLuc reporters containing the stem-loop, there was almost 6-fold increase in luciferase activity that was largely blocked by the combined deletions of the three BFD2 binding sites (Fig. 5A). Individual deletions of BR2 and BR3 showed only a 3-fold induction of luciferase activity by stress (Fig. 5B). The combined BR2+BR3 deletion further lowered induction to 1.8-fold (Fig. 5C), although it is noted that the basal expression of these reporters varied somewhat depending on the presence of BR1. Together, these data suggest that the two BFD2-binding sites most proximal to the *BFD1* CDS (BR2 and BR3) function in an additive fashion to mediate preferential translation of BFD1 in response to stress that induces bradyzoite differentiation.

**Figure 5.**
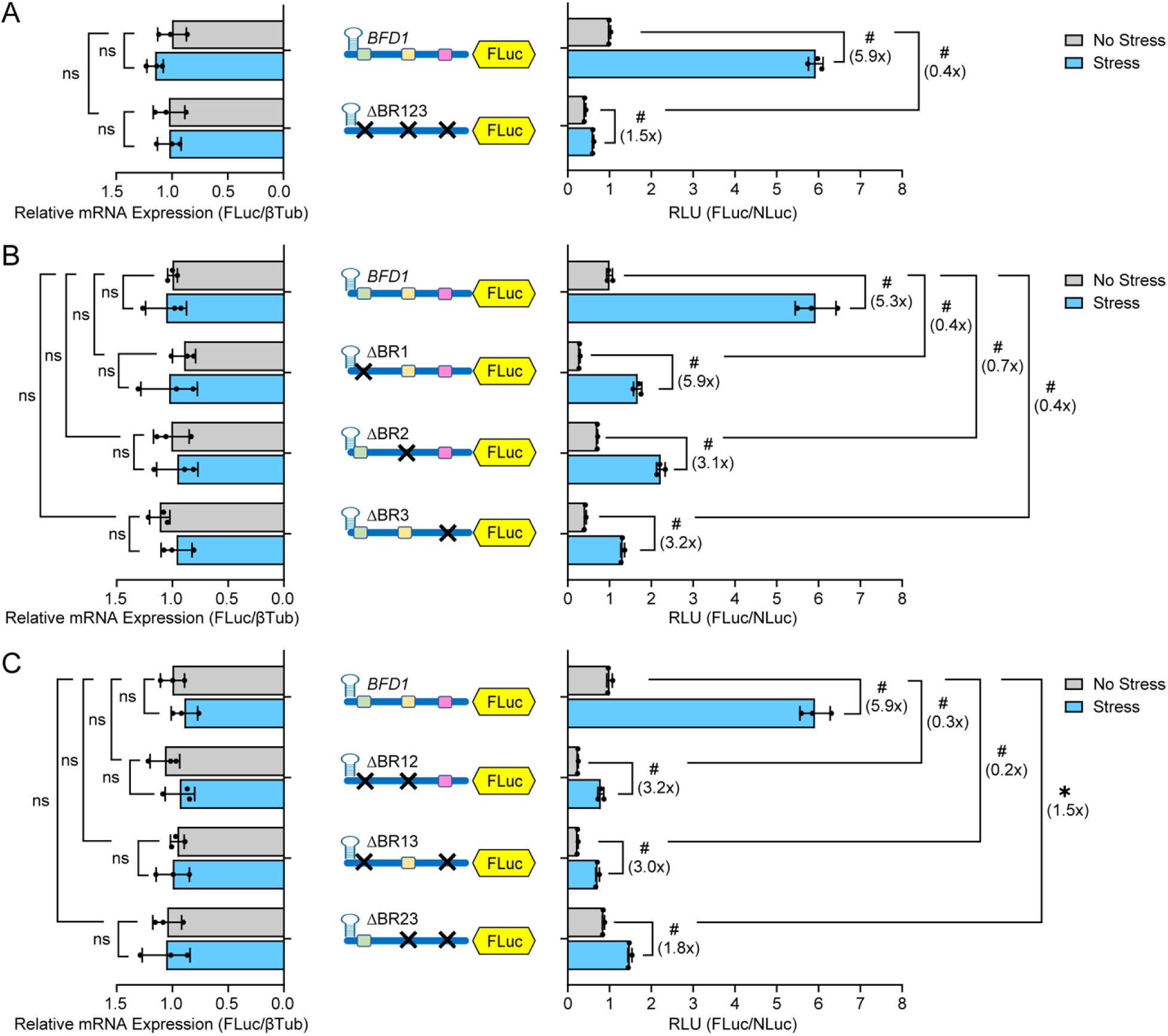
Cap-independent translation of *BFD1* requires BFD2-binding sites. (A-C) 5’-stem-loops were inserted in the BFD1-Fluc reporters with individual deletions (B) or combinations of BFD2 binding sites (A, C), as designed by the “X”. The reporter plasmids reporters were transfected into WT ME49 parasites cultured under alkaline stress (Stress, blue bars) or tachyzoite culture conditions (No Stress, grey bars) for 24 h and relative luciferase activity was measured. Mean fold change ± standard deviation from 3 biological replicates is plotted with normalization to the relative luciferase activity of the WT BFD1-FLuc reporter with no stress. Luciferase reporter mRNAs were measured by RT-qPCR. Mean transcript abundance ± standard deviation from 3 biological replicates is plotted relative to β-Tubulin with normalization to no stress transfected with BFD1-FLuc reporter. ns = p>0.05; * = p≤0.01; # = p≤0.0001 by Student’s two-tailed t-test. Fold changes are shown in parentheses.

### Preferential translation of BFD1 is blunted in parasites lacking *BFD2*

To further determine the role of BFD2 on cap-independent translation of *BFD1* during stress, we replaced the entire *BFD2* CDS with a DHFR-mCherry selectable marker cassette using a dual guide CRISPR/Cas9 strategy (Fig. S6A). Δ*BFD2* clones were validated by PCR of genomic DNA (Fig. S6B). Parasite plaque and replication assays showed no decrease in parasite replication fitness compared to parental ME49 parasites (Fig. S6C-D). However, as gauged by *Dolichos* lectin staining of cyst walls, the Δ*BFD2* parasites showed a robust defect in cyst formation compared to WT ME49 (Fig. S6E-F), consistent with earlier reports (18, 19).

We then transfected our BFD1-FLuc reporter with or without the 5’-stem-loop into WT and Δ*BFD2* parasites. In WT parasites, the BFD1-FLuc again showed ~5-fold increase in preferential translation during stress that was sustained upon insertion of the stem-loop; however, there was only about a ~3-fold increase in luciferase activity in the Δ*BFD2* parasites regardless of the presence of the stem-loop (Fig. 6A). We also measured BFD1-Luc reporters in which the two most critical BFD2-binding sites (BR2+BR3) were removed, with or without the stem-loop included. In both WT as well as Δ*BFD2* parasites, stress-induced preferential translation was curtailed, only increasing ~2-fold regardless of BFD2 presence or the insertion of the stem-loop (Fig. 6B). Collectively, these results further support that BFD2 binding on *BFD1* mRNA is a major contributor in the mechanism driving cap-independent translation of BFD1 in response to stress.

**Figure 6.**
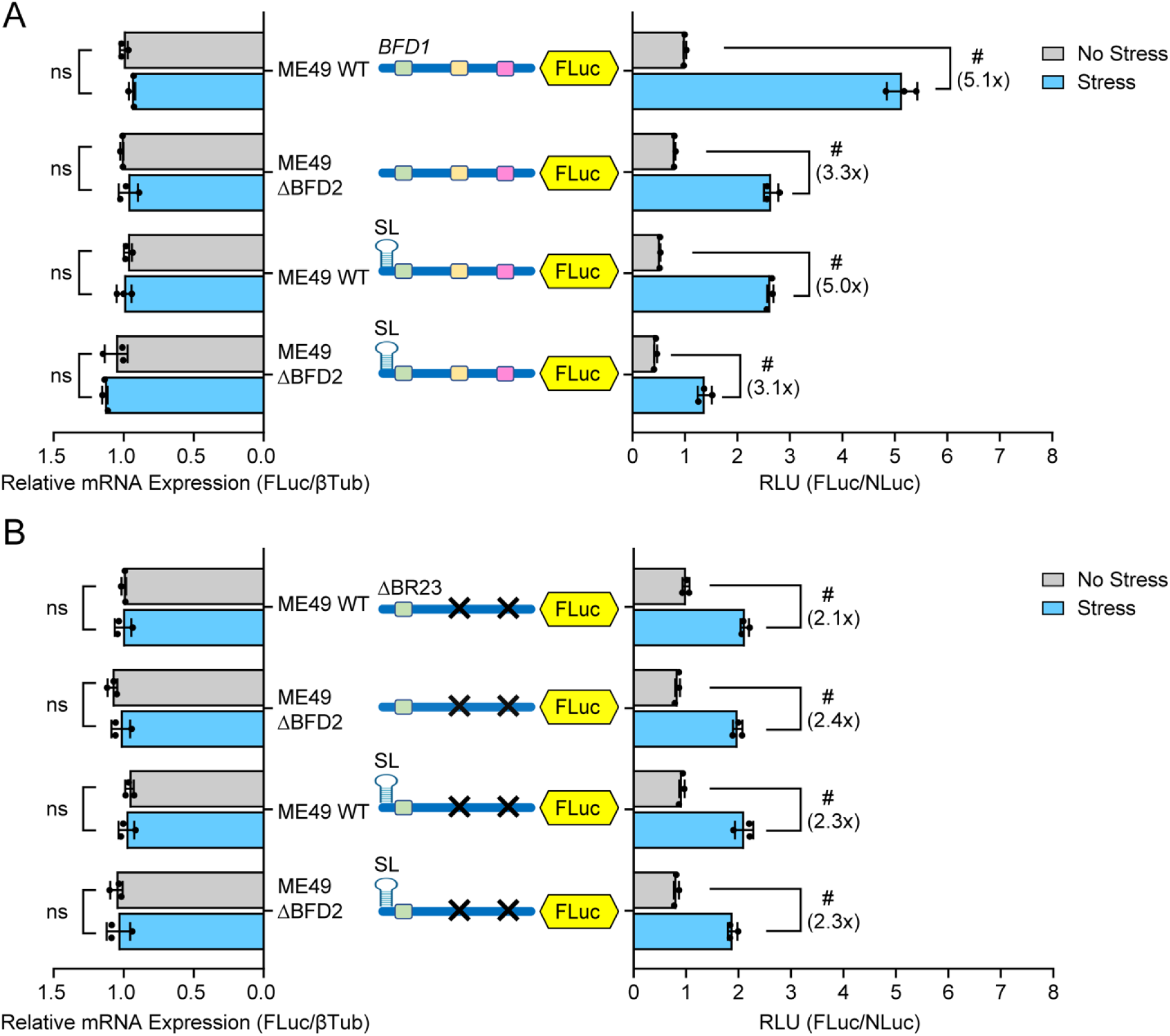
Preferential translation of *BFD1* is blunted in BFD2 knockout parasites. (A) BFD1-FLuc reporters with or without an inserted stem-loop were co-transfected with Tub-NLuc into WT ME49 or ΔBFD2 parasites. Transfected parasites were cultured under alkaline stress (Stress, blue bars) or tachyzoite culture conditions (No Stress, grey bars) for 24 h and RLU were measured. Mean fold change ± standard deviation from 3 biological replicates is plotted with normalization to the RLU of the BFD1-FLuc reporter in ME49 devoid of stress. Luciferase reporter mRNAs were measured by RT-qPCR. Mean transcript abundance ± standard deviation from 3 biological replicates is plotted relative to β-Tubulin with normalization to ME49 tachyzoites transfected with the BFD1-FLuc reporter. ns = p>0.05; # = p≤0.0001 by Student’s two-tailed t-test. Fold changes are shown in parentheses. (B) BFD1-FLuc reporters lacking BFD2-binding sites BR2+BR3 with or without a stem-loop version insertion were co-transfected with Tub-NLuc into WT ME49 or ΔBFD2 parasites and measured and analyzed as described for panel A.

## Discussion

Preferential translation mechanisms are instrumental for *Toxoplasma* conversion of tachyzoites to bradyzoites, which allows the parasite to persist in its host and cause chronic toxoplasmosis (3, 29, 30). In this study, we addressed the processes underlying stress-induced translation of *BFD1*, encoding a transcription factor that is necessary and sufficient for bradyzoite formation (15). The 5’-leader sequence of *BFD1* confers translational control since the native 3’-UTR was absent in our reporter assay, replaced with a heterologous one in our endogenously tagged *BFD1* parasites (Fig. 1). Two independent strategies, stem-loop insertion (Fig. 2) and depletion of cap-binding protein eIF4E1 (Fig. 3), indicated that preferential translation of *BFD1* during stress is driven by a cap-independent mechanism. In contrast, the same strategies showed that another preferentially translated gene *BFD2*, which also contributes to the program of parasite differentiation (18, 19), utilizes a canonical cap-dependent paradigm. We conclude that translational control and differentiation in *Toxoplasma* are directed by cap-independent translation, along with canonical cap-dependent mechanisms.

### Cap-independent mechanisms driving *BFD1* translation during stress

First described for viral RNAs (31, 32), the extent of cap-independent translation of cellular mRNA is still not well understood in eukaryotic cells. Translation of several viral mRNAs that are naturally uncapped is initiated by an internal ribosome entry site (IRES) or cap-independent translation elements (CITEs) (23, 33). Cap-independent recruitment of mRNAs to ribosomes in eukaryotes typically occurs in response to cellular stress or signals to undergo differentiation. Here, we report the importance of cap-independent translation in a protozoal organism subjected to stress-induced differentiation.

The 5’-leader of *BFD1* is unusually long (~2.7 kb) and associates with BFD2 in a stress-dependent manner at three major sites as identified by CLIP-seq (18, 19). As illustrated in the model of *BFD1* translational control presented in Figure 7, our reporter analyses suggest that cap-independent translation of *BFD1* in response to stress that induces differentiation involves BFD2-binding regions designated BR2 and BR3. Enhanced translation of *BFD2* occurs by cap-dependent mechanisms and the ensuing increased levels of BFD2 protein bind to regions of *BFD1* mRNA that likely contain specific sequences and structural features. Along with enhanced expression, there may be post-translational modifications (PTM) of BFD2 and additional associated proteins that also aid in its binding to *BFD1* mRNA and enhanced translation in response to stress. This targeted BFD2 association with regions localized to BR2 and BR3 is suggested to recruit pre-initiating ribosomes for recognition of the *BFD1* start codon and translation of the CDS (Fig. 7).

**Figure 7.**
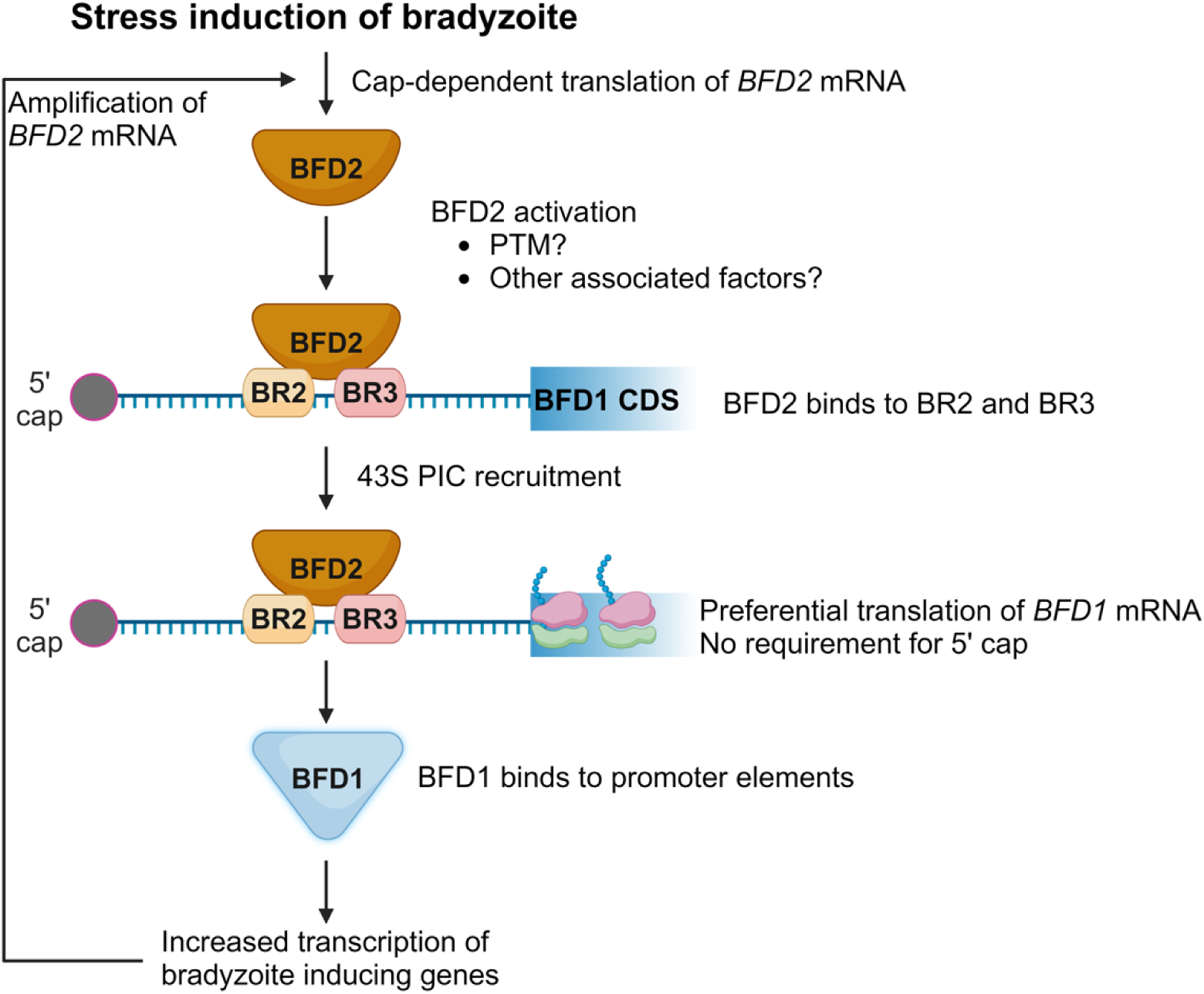
Model for *BFD1* preferential translation during stress-induced differentiation. Cellular stress promotes the cap-dependent preferential translation of *BFD2* mRNA. Increased levels of BFD2 protein contribute to its binding on BR2 and BR3 regions of the *BFD1* 5’-leader. BFD2 PTM and interaction with additional proteins may contribute to its association with the *BFD1* mRNA and preferential translation. BFD2 protein facilitates the translation of the *BFD1* CDS likely by recruiting the preinitiation complex downstream of the 5’-cap and independently of eIF4E1 abundance. BFD1 protein directly binds its target promoters, thus driving the transcriptional response that promotes bradyzoite formation, including increased *BFD2* transcription by positive feedback. Enhanced *BFD2* mRNA contributes to further translation of *BFD2*, amplifying BFD1 expression in response to stress.

It is curious that while the level of translation induction during stress remains upon insertion of the stem-loop at the 5’-end of the *BFD1* mRNA or deletion of the proximal BR1 region, the basal expression and consequently the magnitude of the induced *BFD1* translation is diminished upon stress. A previous report of ribosome profiling in *Toxoplasma* tachyzoites indicates that ribosomes readily translate several 5’-proximal uORFs in the *BFD1* mRNA (26). Given that there are a multitude of *BFD1* predicted uORFs, 37 in total, it is not likely that scanning ribosomes can proceed through the lengthy 5’-leader by leaking scanning or repetitive reinitiation. Rather, the 5’-proximal uORFs could serve as a localized reservoir of ribosomes and associated factors that would become available for stress-induced translation of *BFD1*. Enhanced BFD2 binding to the *BFD1* mRNA during stress would help reposition this reservoir of ribosome pre-initiation complexes (PIC) for appropriate *BFD1* CDS start codon recognition and enhanced translation (Fig. 7). Stress signals trigger increased expression of BFD2 and, along with possible changes in PTMs or associated proteins, may serve as a requisite switch for its engagement with *BFD1* mRNA in this preferential translation mechanism. Increased synthesis of BFD1 would then drive targeted transcription of bradyzoite genes, including *BFD2*, initiating the positive feedback required for cyst formation.

### Preferential translation of *BFD2* occurs by cap-dependent processes

Preferential translation of *BFD2* is also suggested to occur in response to stress but this process occurs by canonical cap-dependent mechanisms (Figs. 2, 3, and 7). The 5’-leader sequences of *BFD2* mRNA also contain predicted uORFs that may confer translational control during stress-induced differentiation. In other model systems, uORFs are integral for cap-dependent preferential translation by processes involving stress induced phosphorylation of the α subunit of eIF2 (17). In these preferentially translated mRNAs with multiple uORFs, the phosphorylated eIF2α directs delayed translation reinitiation that allows for bypass of inhibitory uORFs and enhanced CDS translation (17). Upon stresses that induce bradyzoite development, we reported rapid phosphorylation of *Toxoplasma* eIF2α (TgIF2α) (29). Phosphorylated TgIF2α may contribute to enhanced *BFD2* cap-dependent translation, consequently binding to BFD1 mRNA and promoting its cap-independent preferential translation. In addition to *BFD1* mRNA, BFD2 binds a multitude of other mRNAs, including sequences at the 5’-leaders and 3’-UTRs (19), so the switch may be amplified as the BFD1-directed transcriptome is implemented.

The discovery that two stress-dependent factors crucial to bradyzoite differentiation are preferentially translated with different reliance on the 5’-cap and associated eIF4E1 provides the basis for future interrogation of diverse translational control mechanisms in *Toxoplasma*. Our results can explain why we recently found that the cap-binding protein eIF4E1 is downregulated during stress-induced bradyzoite differentiation in *Toxoplasma* (26). Loss of eIF4E1 would decrease cap-dependent translation, making initiation factors more available for interaction with mRNAs possessing features for cap-independent translation. In closing, our findings represent the first evidence of cap-independent translation associated with stress and differentiation in a protozoan, highlighting the antiquity of this mechanism of translational control in eukaryotic cell evolution. Pharmacological manipulation of translational control mechanisms in pathogenic protozoa may thwart the ability of these parasites to maintain latent forms or reactivate into acute infection.

### Experimental Procedures

#### Host cell and parasite culture

Human foreskin fibroblast (HFF) (ATCC, SCRC-1041) cells were cultured in Dulbecco’s modified Eagle medium (DMEM) (GIBCO, 10-017-CM) pH 7.4, supplemented with 10% inactivated fetal bovine serum (FBS) (R&D systems, S11150), in a humidified 5% CO_2_ incubator at 37°C. WT ME49 strain parasites were provided by Dr. Vern Carruthers (University of Michigan). ME49 tachyzoites were cultured in DMEM supplemented with 5% inactivated FBS. Alkaline medium for stress conditions contained RPMI 1640 (GIBCO, 31800-022) medium supplemented with 5% FBS and 50 mM HEPES. Alkaline medium was adjusted to pH 8.2 using NaOH. Parasites in alkaline medium were grown in ambient CO_2_ and the medium was changed daily.

#### Parasite growth assays

Intracellular ME49WT and ΔBFD2 parasites were mechanically harvested by syringe passaging. 1000 parasites were used to infect confluent HFF cells grown on plates in tachyzoite medium. The plate was then incubated undisturbed for 14 days until plaques were formed. To stain the monolayer, infected cells were rinsed with PBS, fixed with methanol for 15 minutes, then treated with a crystal violet solution for 30 minutes followed by washing with distilled water. The plates were air-dried, and plaques were counted.

For replication assays, purified parasites were used to infect HFF monolayers and allowed to invade for 8 h in tachyzoite medium. After invasion, the medium was replaced to remove uninvaded parasites. After 16 h, the number of parasites per vacuole was counted.

#### Generation of parasites expressing endogenously tagged BFD1

All primers used in this study are listed in Table S1 and were procured from Integrated DNA Technologies. The *BFD1* genomic locus (TGME49_200385) (15) was tagged with three C-terminal hemagglutinin (3xHA) epitope tags in ME49 at the endogenous *BFD1* locus using CRISPR/Cas9-based and double homologous recombination (designated BFD1^HA^). A single guide RNA (sgRNA) sequence near the stop codon of BFD1 was chosen using the EuPaGDT tool (http://grna.ctegd.uga.edu/) and the sequence was cloned into a pCas9-GFP plasmid (34) by site directed mutagenesis using PCR, designated as pCas9-BFD1. Parasites were co-transfected with pCas9-BFD1 along with a PCR amplicon repair template using a Lonza Biosciences Nucleofector. The repair template encoded a 3xHA tag sequence and a dihydrofolate reductase-thymidylate synthase (DHFR*-TS) selectable marker flanked by 40bp homologous to either site of the CAS9 cut site to facilitate homology-directed repair of the CAS9-induced break. The parasites were selected for DHFR*-TS integration by three passages in medium containing 1.0 μM pyrimethamine (Sigma, SML3579) and cloned by limiting dilution. The modified locus of the transgenic clone was validated by PCR analyses of both 5’-end and 3’-end.

#### Generation of FLuc knockout in eIF4E1^mAID-HA^ parasites

A Cas9 plasmid encoding a single guide RNA directed to the FLuc CDS, termed pCas9-FLuc, was co-transfected along with a PCR amplicon of a genomic HXGPRT copy into ME49 eIF4E1^mAID-HA^ parasites, which express an integrated copy of FLuc and lack HXGPRT using a Lonza Biosciences Nucleofector. The parasites were selected for three passages in medium containing 25 µg/mL mycophenolic acid (Sigma, M5255) and 50 µg/mL xanthine (Sigma, X7375) and cloned by serial dilution. The knockout clones were confirmed by measuring firefly luciferase signal for 2×10^6^ parasites.

#### Generation of *BFD2* knockout parasites (Δ*BFD2*)

Two gRNAs were designed to replace the entire *BFD2* coding in ME49 parasites using a strategy that has previously been outlined (35). Both gRNAs were cloned separately in pCas9-GFP plasmid to target the 5’- and 3’-end of the *BFD2* coding sequence, designated as pCas9-gRNA1 and pCas9-gRNA2. U6-gRNA1 region was amplified and digested with Kpn I and Xho I restriction enzymes. The resulting DNA insert was ligated into the vector pCas9-gRNA2 digested with the same enzymes using T4 DNA ligase. The resulting plasmid is pCas9-dualgRNA. A double-stranded donor containing DHFR-mCherry flanked by 40 bp segment homologous to the 5’-leader of *BFD2* and a 40 bp region complementary to the 3’-UTR regions of the *BFD2* gene was generated by PCR-amplification. Co-transfection of pCas9-dualgRNA and the DHFR-mCherry repair template into ME49 parasites was performed using a Nucleofector (Lonza Biosciences). The parasites were selected for three passages in medium containing 1.0 μM pyrimethamine (Sigma, SML3579) and cloned by limiting dilution. The *ΔBFD2* clone was validated by PCR analyses.

### Immunofluorescence assays

HFF cells were grown to confluency on coverslips and infected with WT ME49 or BFD1^HA^ tachyzoites. Four hours post-invasion, the medium was changed to alkaline medium and incubated for 24 h. Coverslips were fixed with 4% paraformaldehyde (Sigma, P6148) and washed with PBS. Infected cell monolayers were then blocked and permeabilized with blocking buffer containing 3% bovine serum albumin (BSA) (Sigma, A9418) and 0.2% Triton X-100 (Sigma, 93443) in PBS. A rat anti-HA antibody (Roche, 11867423001) at 1:1000 dilution in blocking buffer was applied was applied to the coverslips at 4°C for 16 h. After washing, the coverslips were incubated with secondary antibody, Alexa Fluor 598 anti-rat (Thermo Fisher, A-11007) diluted at 1:5000 in blocking buffer and 1 μg/ml DAPI (Invitrogen, D1306) for 1 h at room temperature in the dark. The coverslips were mounted in ProLong gold antifade mounting solution (Invitrogen, P36930) and examined by fluorescent microscopy.

For the bradyzoite differentiation assay, WT ME49 and ΔBFD2 parasites were cultured on separate coverslips under bradyzoite conditions for 3 days. The IFA was conducted as detailed above. Fluorescein isothiocyanate (FITC)-conjugated *Dolichos biflorus* lectin (DBL) (Vector Laboratories, FL1031) at 1:300 dilution was included along with the secondary antibody to reveal the cyst wall.

### Immunoblot analyses

Parasites grown in confluent HFF cells under unstressed and alkaline stress culture conditions for 24 h or 5 days were lysed in RIPA buffer (25 mM Tris pH 7.4, 150 mM NaCl, 0.1% SDS, 1% NP-40, 0.5% sodium deoxycholate) supplemented with a protease and phosphatase inhibitor cocktail (Thermo Fisher, 78425). The lysed samples were sonicated on ice followed by centrifugation at 13,000 xg. The supernatant was boiled for 10 min in 1X SDS-PAGE protein sample loading buffer (Invitrogen, NP0008) supplemented with 10% β-mercaptoethanol (Sigma, M3148). The samples were then separated by electrophoresis using a 4% to 12% Tris-Bis polyacrylamide gradient gel (Invitrogen, NP0326BOX) and transferred to a pre-equilibrated nitrocellulose membrane (Cytiva, 10600001). The membrane blots were blocked in blocking buffer containing 5% non-fat milk/TBST (20 mM Tris, 150 mM NaCl, 0.1% (w/v) Tween 20) for 1 h at room temperature. Primary antibody was applied at 4°C for 16 h using rat anti-HA (Roche, 11867423001) for HA-tagged proteins and rabbit anti-BAG1 (provided by Dr. Vern Carruthers, University of Michigan) polyclonal antibodies at 1:1000 dilution, rabbit anti-tubulin (provided by Dr. David Sibley, Washington University), or rabbit anti-aldolase (provided by Dr. David Sibley, Washington University) polyclonal antibodies at 1:5,000 dilution in blocking buffer. Secondary antibody was applied using appropriate HRP-conjugated antibodies (GE healthcare) at a 1:5,000 dilution in blocking buffer. Immunoblots were imaged using SuperSignal West Femto reagent (ThermoFisher, 34095) and chemiluminescence was detected via a Bio-Rad ChemiDoc Imaging System.

### Reporter plasmid construction

The promoter and 5’-leader sequence of the *tubA1* gene were PCR-amplified from ME49 genomic DNA. The promoter and transcriptional start sites were determined using RAMPAGEseq and H3K9ac ChIP-chip data available on toxodb.org (25, 36, 37). The firefly luciferase CDS was PCR-amplified from pGL3 plasmid (Promega). The PCR products were ligated using In-Fusion cloning kit (Takara, 638947) and the resultant plasmid was named Tub-FLuc. To generate the BFD1 reporter construct, the *TUBA1* 5’-leader sequence was replaced by the *BFD1* 5’-leader and the first 35 codons of the *BFD1* CDS to maintain an out-of-frame uORF. The BFD2 reporter construct was generated by replacing the *tubA1* 5’-leader with that of the short *BFD2* leader as determined by its eIF4E1-binding sites (26) (Fig S3B, GSE243203). Each of these FLuc reporters included the Tub promoter, the respective encoded 5’-leaders and the FLuc CDS. For the transfection control reporter construct, the firefly luciferase CDS was replaced by Nano luciferase from pNL1.1 (Promega) using In-Fusion cloning. The resulting plasmid Tub-NLuc features the Tub gene promoter and encoded 5’-leader juxtaposed to the Nano CDS.

The sequence of the 5’-stem-loop, obtained from a previous study (28), was inserted into relevant constructs by PCR and plasmids were generated using the KLD Enzyme Mix (New England Biolabs, M0554S). Similarly, single deletions of BFD2 Binding Regions (BR) 1-3 were performed by PCR followed by a KLD reaction. The regions that were deleted were selected based on previously published CLIPseq data (19) (GSE211957). Region BR1 corresponds to 144-354 nt (211 nt total) of the *BFD1* mRNA leader; BR2 corresponds to 1431-1722 nt (292 nt total); BR3 corresponds to 2470-2655 nt (186 nt total). Combinations BR1, 2 and 3 deletions were obtained by PCR paired with KDL reaction by using single BR deletion plasmids as a template. All plasmids were verified by Sanger sequencing over the length of their 5’-leaders. Primers used for constructing plasmids are listed in Table S1.

### Reporter assays

Luciferase reporters along with Tub-Nano construct were co-transfected into 2×10^6^ ME49 parasites using a Lonza Biosciences Nucleofector and allowed to infect confluent HFFs in 6-well plates. Four h post-invasion, the medium was replaced with normal pH media (No Stress) or alkaline pH (8.2) medium (Stress) and incubated for 24 h at 37°C. After washing in PBS, 500 μl of 1× passive lysis buffer solution (Promega, E1941) was added to each well. Plates were placed on an orbital shaker at 100 rpm for 10 min at room temperature to ensure cell lysis. Cell lysates were then clarified by centrifugation. 20 μl of the supernatant was used for measuring FLuc and NLuc values according to manufacturer’s protocol (Promega, N1630) using a 20/20^n^ luminometer (Turner Biosystems) and the remainder of the lysates were used for qPCR measurements. The FLuc and NLuc values were determined, and the FLuc to NLuc ratios (Relative Luciferase Units - RLU) were calculated for each. The RLU of transfected parasites with the respective control plasmid grown in unstressed conditions was adjusted to 1.0. The fold-change due to alkaline stress was calculated by RLU under stress conditions normalized to the RLU of unstressed condition. The fold change in luciferase activity between two different constructs was measured by one reporter’s RLU under no stress conditions normalized to the RLU of control reporter.

### Real-time reverse transcription PCR (qRT-PCR)

Total RNA was isolated from the infected cell lysates using TRIzol-chloroform according to manufacturer’s protocol (Invitrogen, 10296028). cDNA synthesis was performed using SuperScript IV First-Strand cDNA synthesis system (Invitrogen, 18091300) according to the manufacturer’s protocol. cDNA was diluted 1:10 for BFD1^HA^ parasites and 1:5 for reporter samples and used for real-time PCR with SYBR Green PCR Master Mix (Applied Biosystems, 4368708) reagent using an Applied Biosystems QuantStudio 5 Real-Time PCR system. *Toxoplasma* β-tubulin (TGME49_266960) was used for normalization. We verified that GAPDH (TGME49_289690) mRNA levels did not change under alkaline stress conditions when β-tubulin mRNA was used for normalization. All primers used in qPCRs are listed in Table S1.

## Data availability

Data presented in the manuscript, plasmids, and other reagents are available for academic purposes upon request.

## Supporting information

This article contains supporting information, including tables and figures in a single file.

## Acknowledgments

The authors thank Drs. Vern Carruthers and David Sibley for kindly providing reagents.

## Author contributions

V.D., M.J.H., R.C.W., and W.J.S. conceptualization; V.D., M.J.H., M.S.B., and W.J.S. data curation; V.D., M.J.H., M.S.B, R.C.W., and W.J.S. formal analysis; R.C.W. and W.J.S. funding acquisition; V.D., M.J.H., M.S.B., R.C.W., and W.J.S. investigation; V.D., M.J.H., R.C.W., and W.J.S. methodology; R.C.W., and W.J.S. project administration; R.C.W., and W.J.S. resources; R.C.W., and W.J.S. supervision; V.D., M.J.H., M.S.B., R.C.W., and W.J.S. visualization; V.D., M.J.H., R.C.W., and W.J.S. writing–original draft; V.D., M.J.H., R.C.W., and W.J.S. writing–review and editing.

## Funding and additional information

This work was supported by the National Institutes of Health (AI172752 and AI167662 to W.J.S. and R.C.W.) and the Showalter Trust (W.J.S. and R.C.W.).

---

**Supplementary Figure 1.**
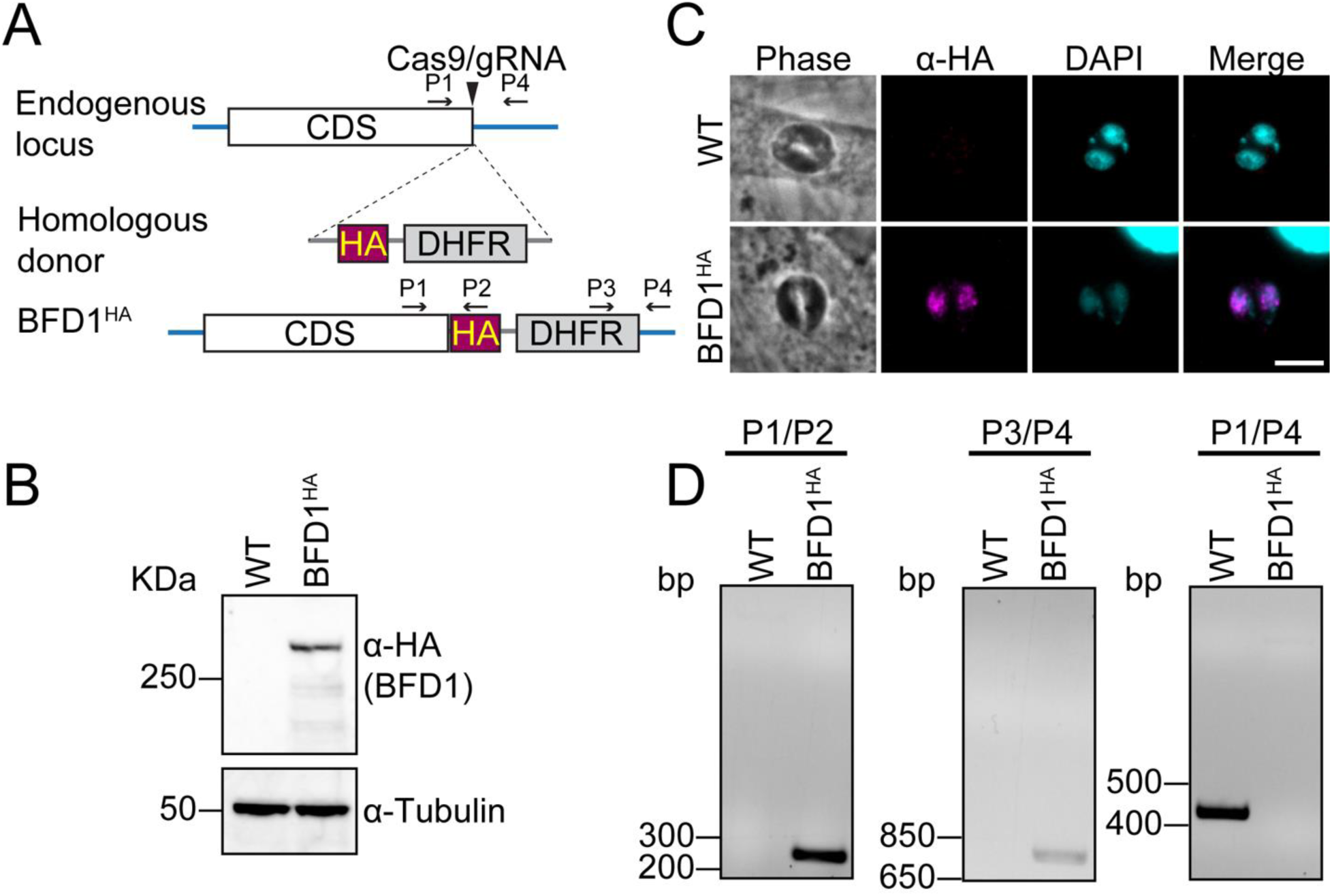
Endogenous tagging of BFD1^HA^. (A) Schematic of the CRISPR/Cas9 approach to fuse endogenous BFD1 with a C-terminal HA epitope tag in ME49 strain parasites. DHFR served as a selectable marker. (B) Immunoblot of WT (parental) ME49 or BFD1^HA^ lysates after 24 h alkaline stress probed with anti-HA. Anti-tubulin was probed as a loading control. (C) IFA of WT ME49 and BFD1^HA^ parasites after 24 h alkaline stress using anti-HA (magenta). DAPI (blue) was used to visualize nuclei. Scale bar = 5 μm. (D) Diagnostic PCRs of gDNA were used to validate the proper integration of the HA tag using primers shown in (A).

**Supplementary Figure 2.**
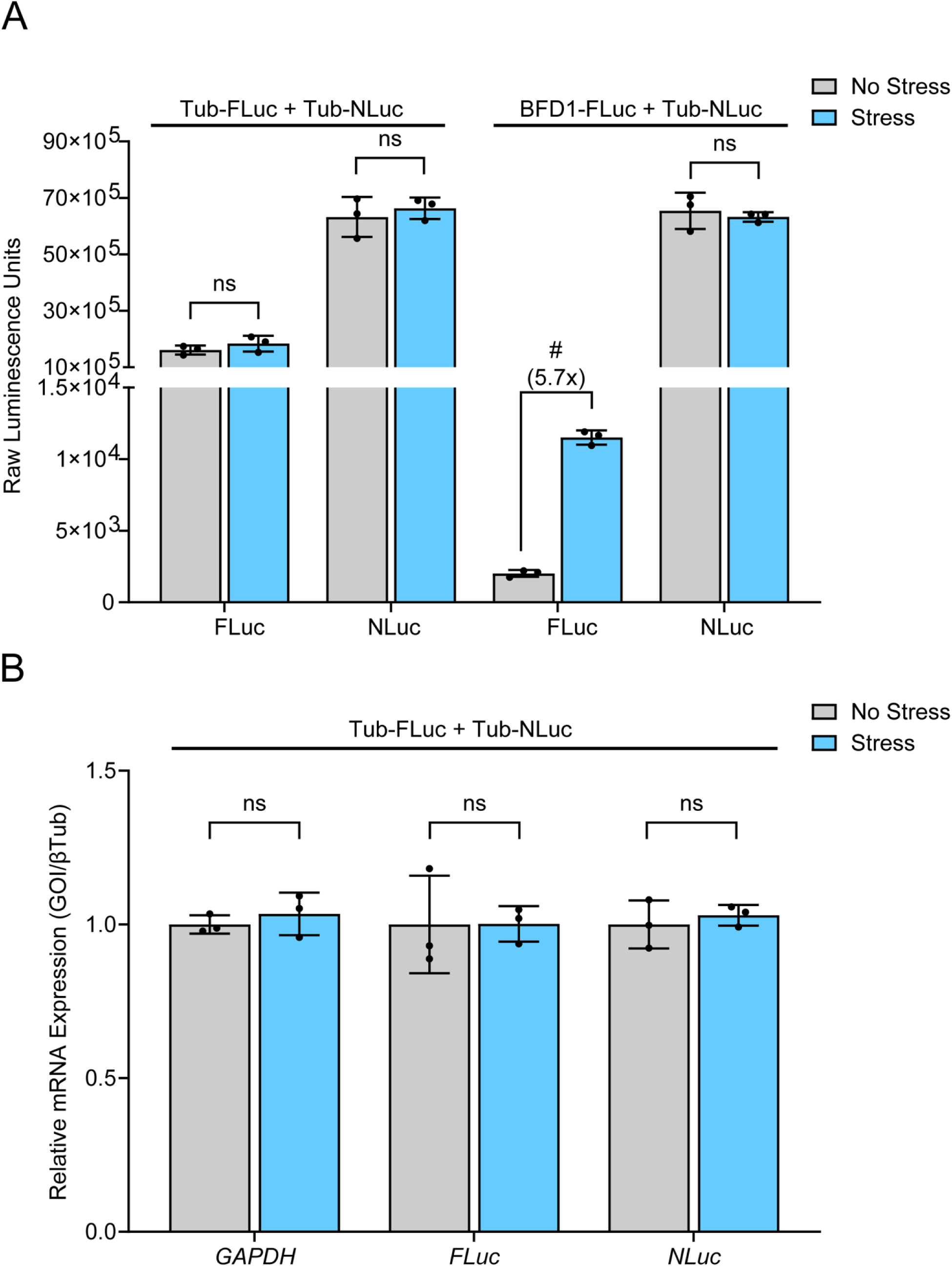
Alkaline stress does not affect NLuc activity or its mRNA levels. The raw values of FLuc and NLuc measurements used to generate Fig. 1C are presented. (A) Luciferase activity in alkaline stress (Stress, blue bars) or tachyzoite conditions (No Stress, grey bars). Mean luciferase activity ± standard deviation from 3 biological replicates is plotted. ns = p>0.05; # = p≤0.0001 by Student’s two-tailed t-test. (B), Mean transcript abundance ± standard deviation from 3 biological replicates plotted relative to β-Tubulin with normalization to tachyzoites. GAPDH is included as an additional normalization to further support that β-Tubulin levels do not change in response to stress. ns = p>0.05 by Student’s two-tailed t-test. Fold changes are shown in parentheses.

**Supplementary Figure 3.**
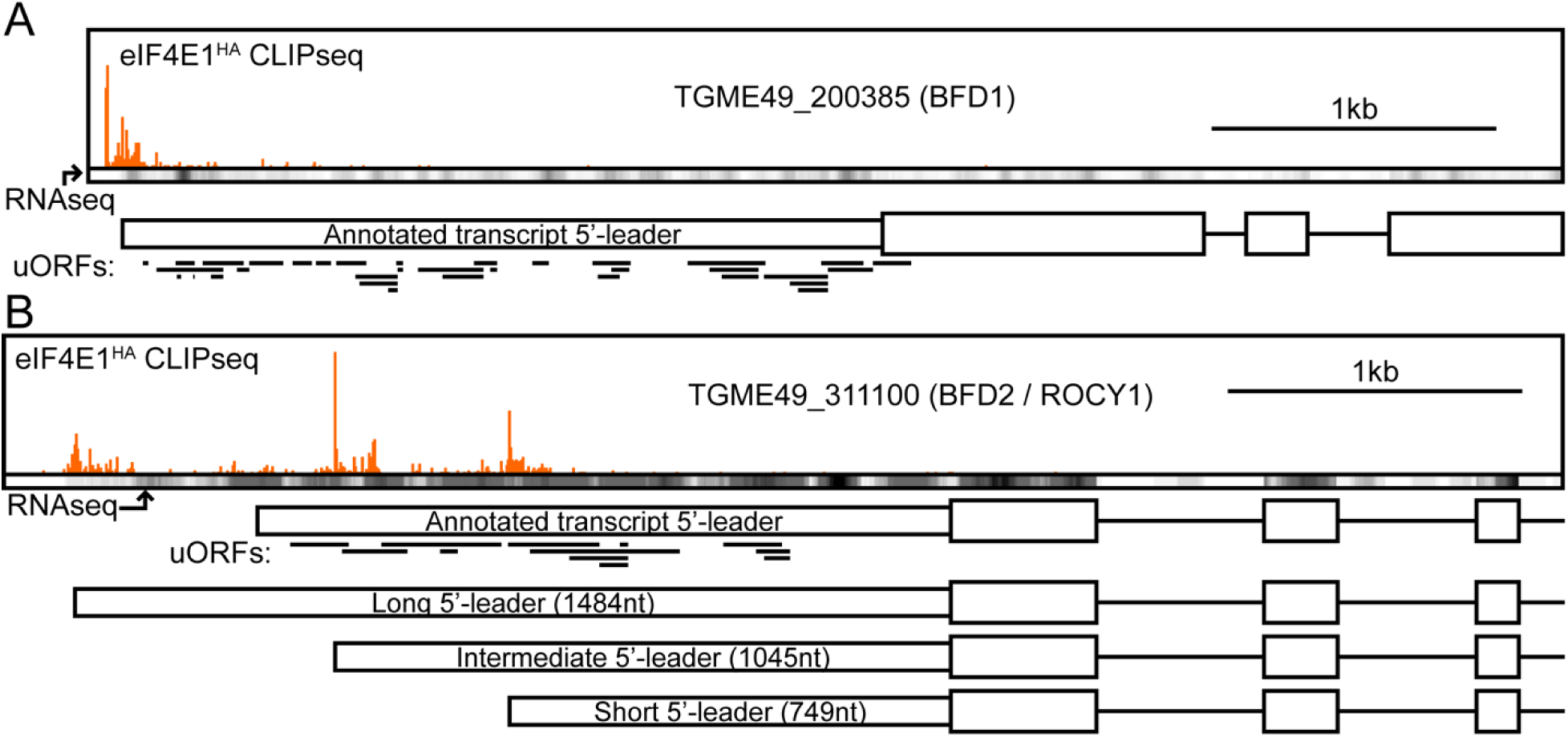
Transcriptional start site predictions of *BFD1* and *BFD2* transcripts. eIF4E1-mRNA interaction sites as evidenced by CLIPseq are displayed as orange profiles above the gene models for *BFD1* (A) and *BFD2* (B). CLIPseq data was obtained from GSE243203. Strand-specific RNAseq heatmaps, obtained from GSE243206, are displayed below CLIPseq data to support transcript abundance over the gene bodies. 5’-leaders are displayed as narrow boxes, CDS as thick boxes, introns as lines. Predicted uORFs are displayed as thick black lines below the gene model. (A) The data presented is consistent with a single transcriptional start site corresponding to the annotated gene model for *BFD1*. (B) The data presented is inconsistent with the annotated *BFD2* 5’-leader and instead supports three transcript isoforms generated by alternate transcriptional start site usage, with a major 5’-leader ~750 nt in length used in the BFD2-FLuc reporter assays.

**Supplementary Figure 4.**
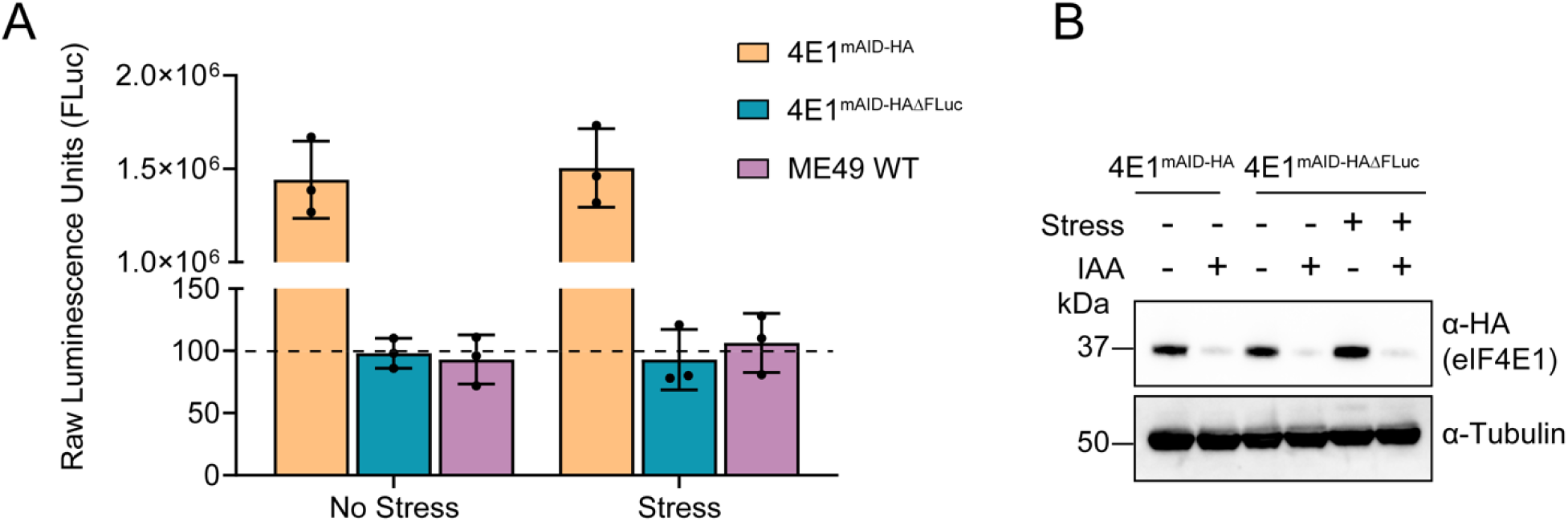
Knockout of FLuc in eIF4E1^mAID-HA^ parasites. (A) Raw FLuc activity was measured for 10^6^ eIF4E1^mAID-HA^ parasites containing integrated FLuc and eIF4E1^mAID-HA^**^Δ^**^FLuc^ parasites in which the integrated FLuc gene was knocked out, under stress or unstressed conditions. The dotted line indicates the limit of luciferase activity detection in WT ME49 parasites as determined by taking the average from three independent measurements of untransfected parasites. (B) Immunoblot analyses with anti-HA to measure depletion of eIF4E1^mAID-HA^ in response to 4 h treatment with IAA in stress or unstressed conditions. Tubulin was probed as a loading control. Molecular weight markers are shown in kDa.

**Supplementary Figure 5.**
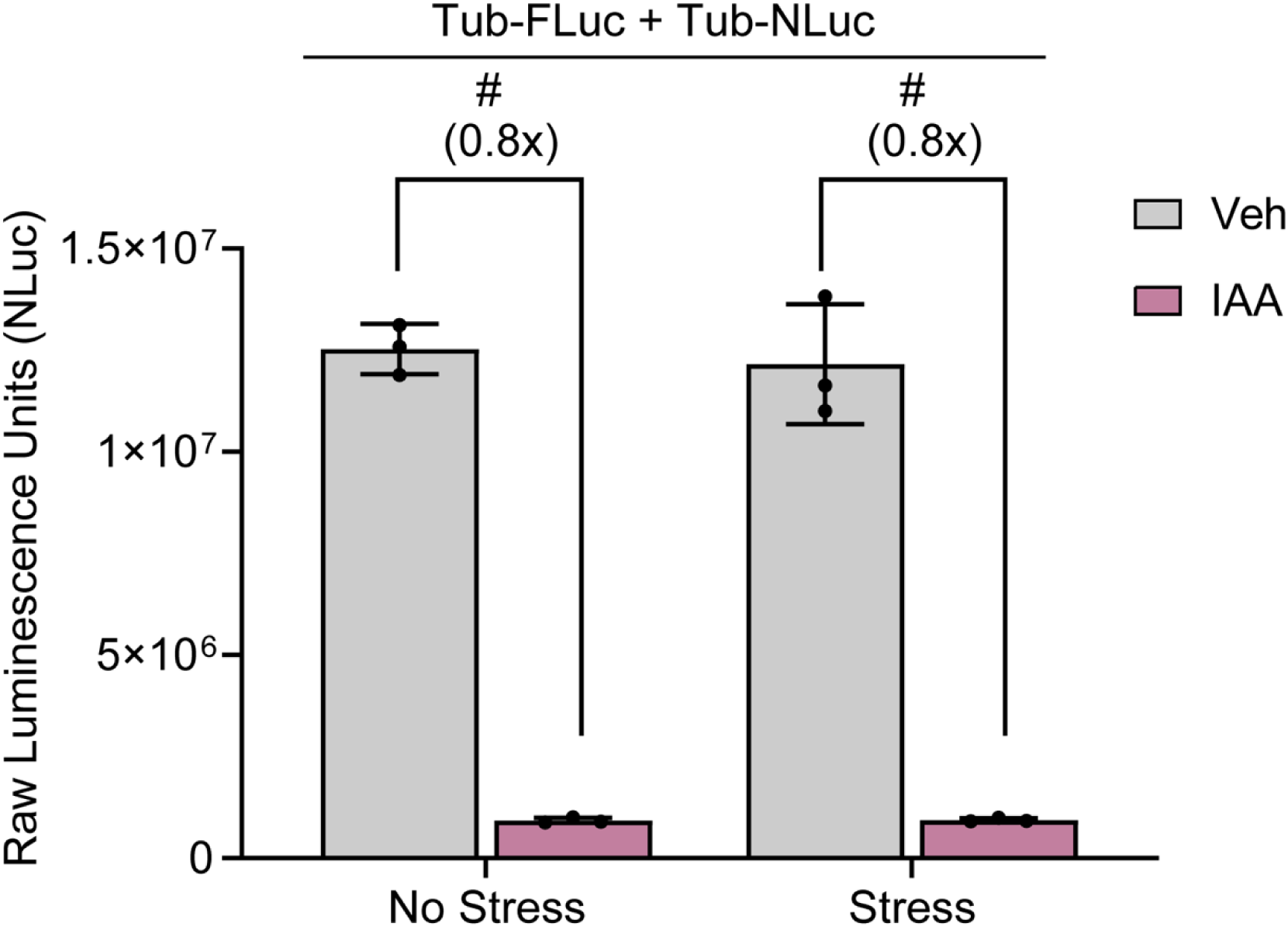
NLuc activity decreases in response to eIF4E1 depletion. eIF4E1^mAID-HAΔFLuc^ parasites were co-transfected with Tub-FLuc and Tub-NLuc. Raw values of NLuc activity are shown. Bar graph represents mean NLuc activity ± standard deviation from 3 biological replicates. # = p≤0.0001 by Student’s two-tailed t-test. Fold changes are shown in parentheses.

**Supplementary Figure 6.**
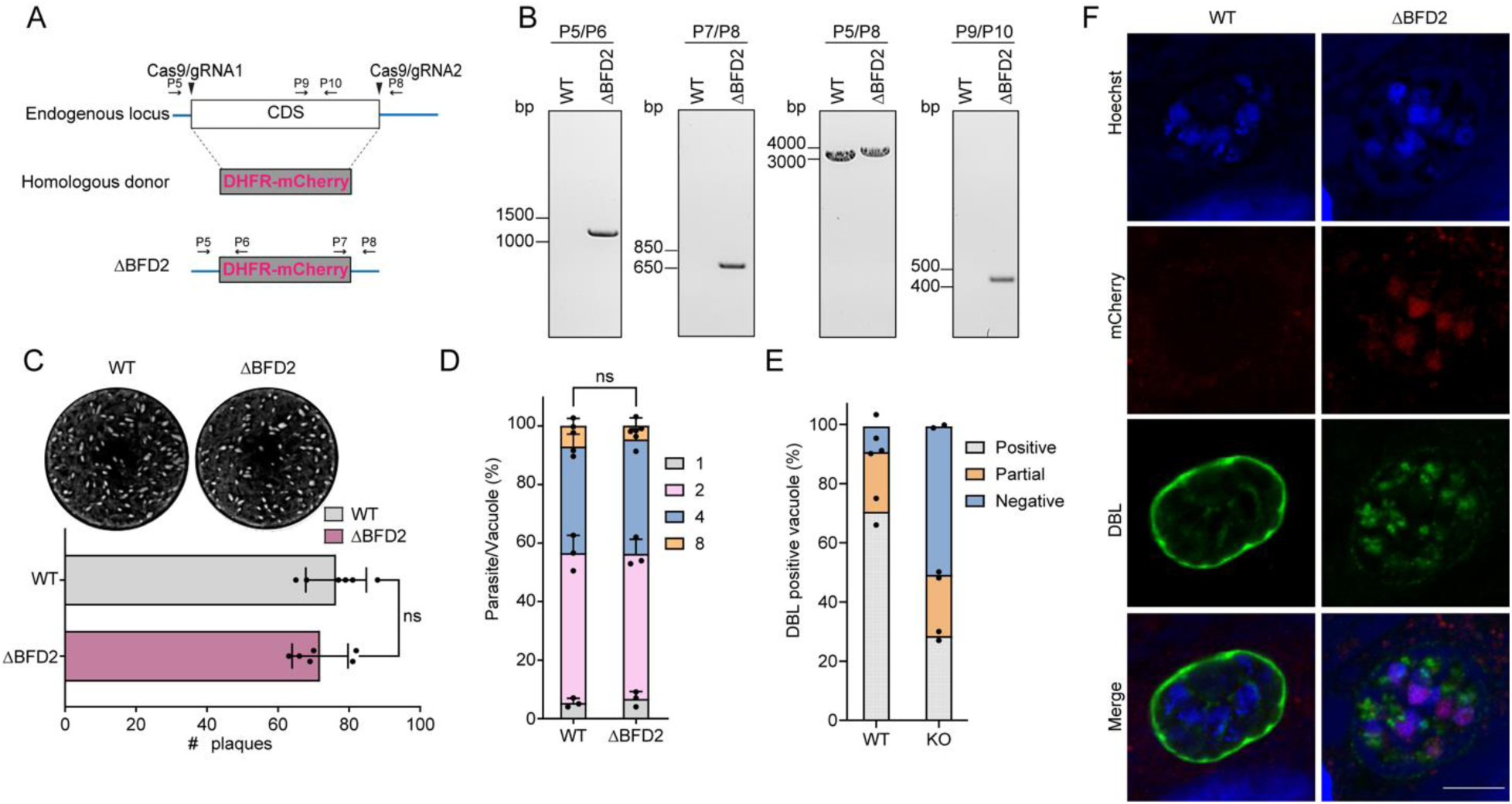
Generation and phenotyping of BFD2 knockout parasites. (A) Schematic of CRISPR/Cas9 approach to generate ΔBFD2 parasites. DHFR-mCherry fusion operates as a selectable marker. (B) Genomic PCRs using primers shown in (A) that confirm replacement of the *BFD2* CDS with the DHFR-mCherry cassette. (C) Representative plaque assay of WT ME49 or ΔBFD2 parasites grown under tachyzoite culture conditions for 14 days. Quantitation of plaque number is included. ns = p>0.05 by Student’s two-tailed t-test. (D) Replication assay showing number of parasites per vacuole in WT ME49 and ΔBFD2 parasites 16 h post-invasion. Mean number of parasites per vacuole ± standard deviation is plotted from 3 biological replicates, with a minimum of 100 random vacuoles counted per sample. (E) Quantitation of parasite differentiation of WT ME49 and ΔBFD2 parasites after 72 h of alkaline stress from two biological replicates. Cultures were stained with FITC-labeled *Dolichos biflorus* lectin (DBL) to visualize cyst walls. A minimum of 100 random vacuoles were counted per sample. (F) Representative vacuoles counted in (E) after 72 h alkaline stress. DBL (green) stains cyst wall. mCherry (red) exclusively stains ΔBFD2 parasites. Hoechst (blue) used to visualize nuclei. Scale bar = 10 μm.

**Supplemental Table S1.**
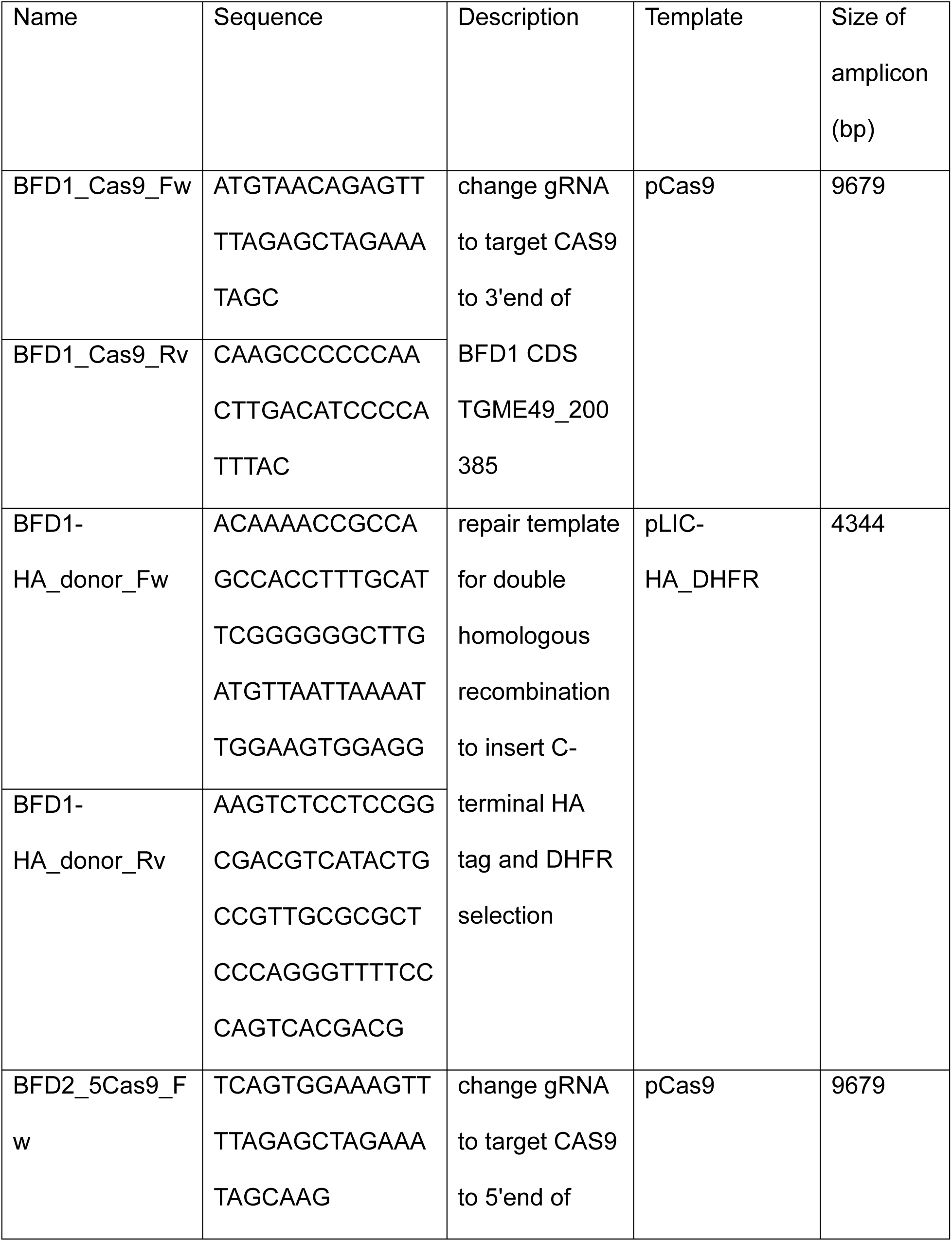
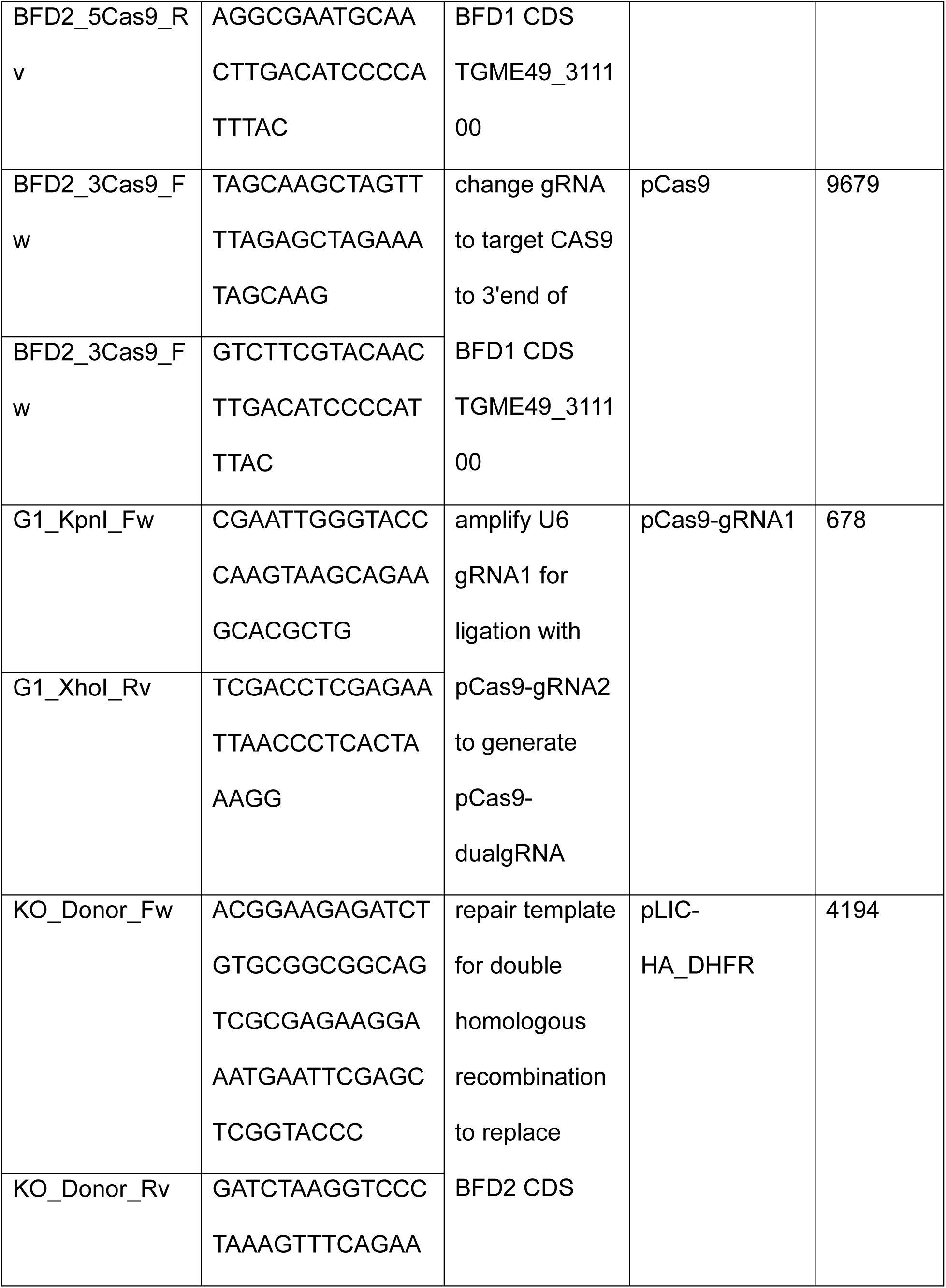
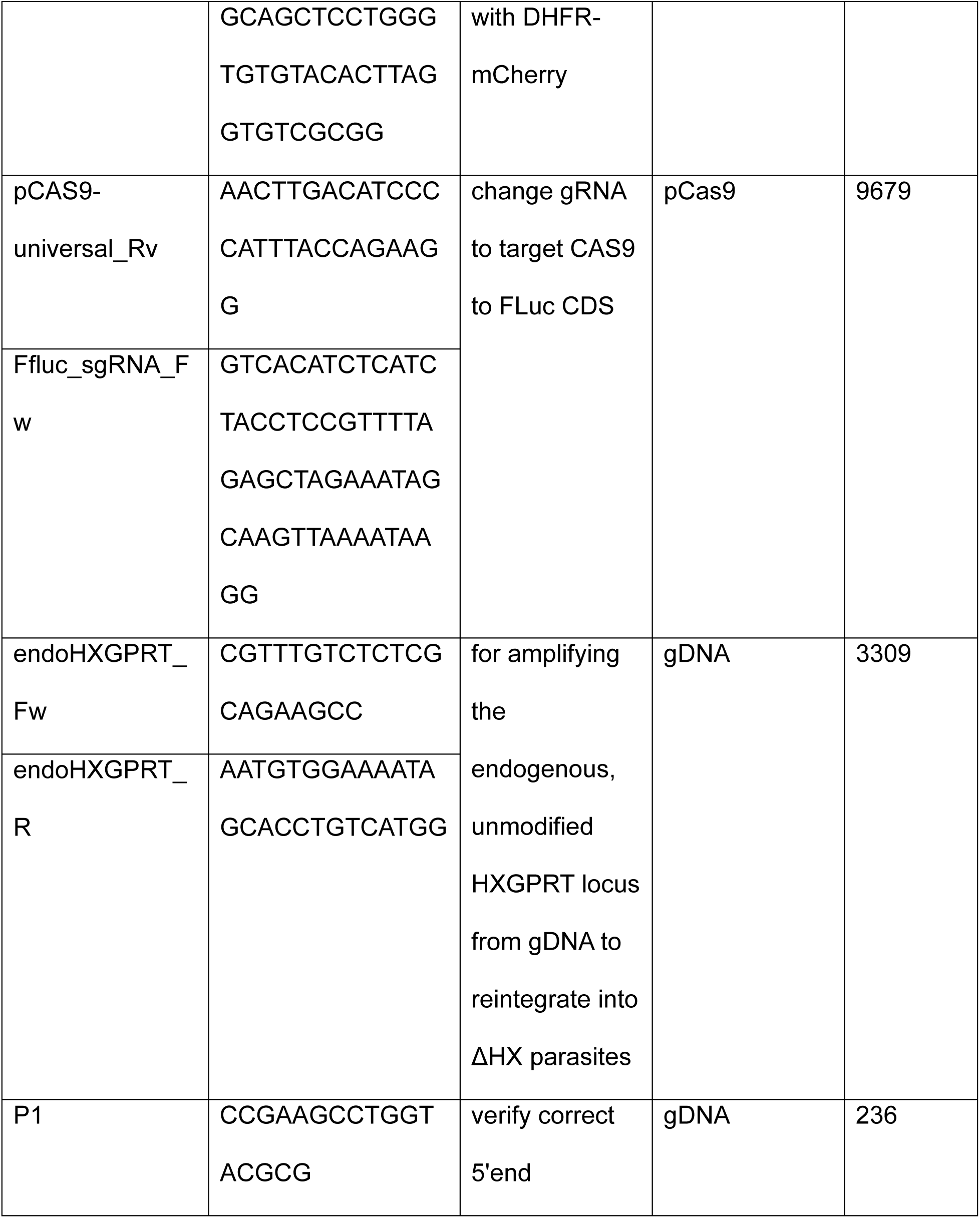
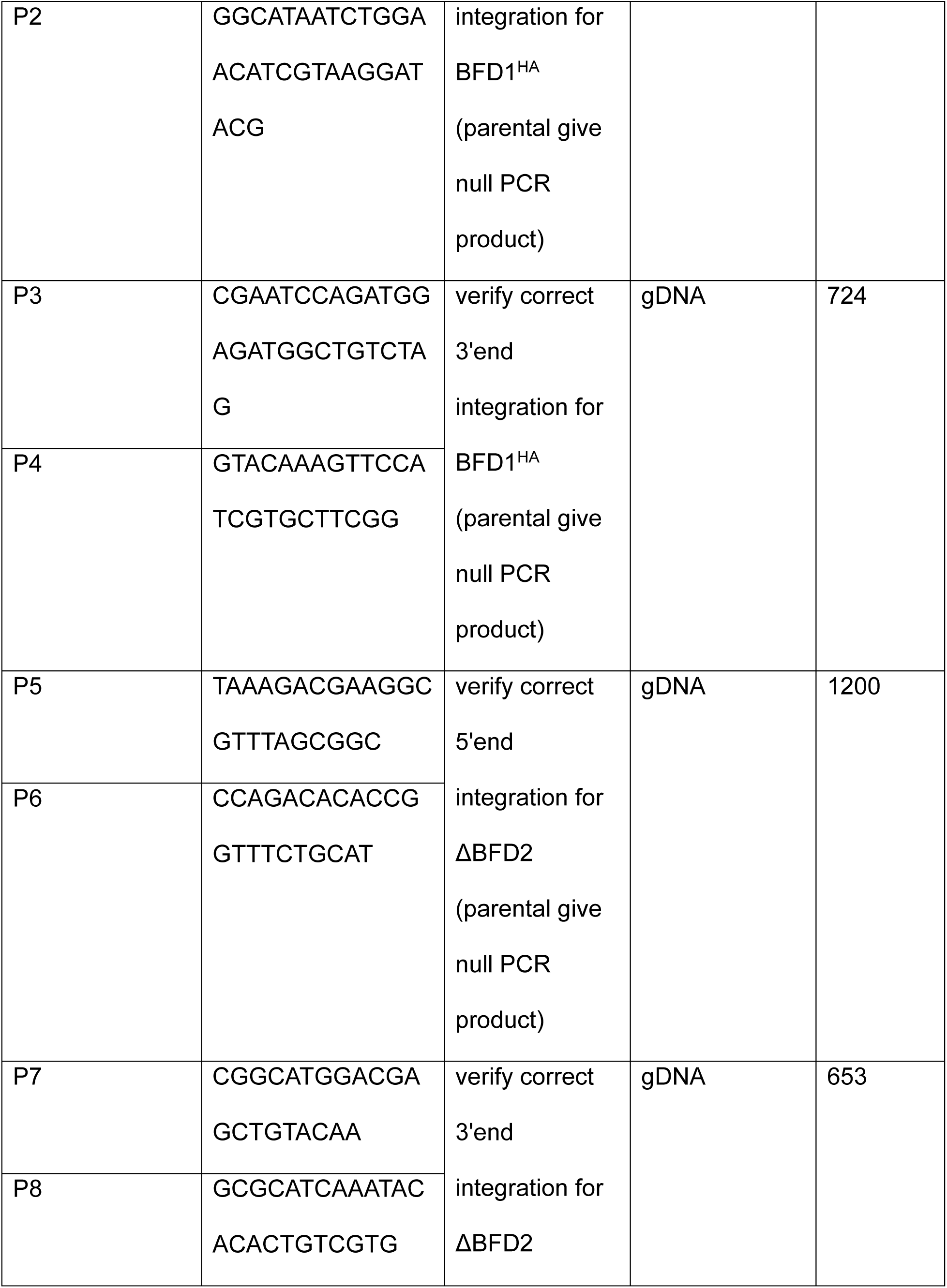
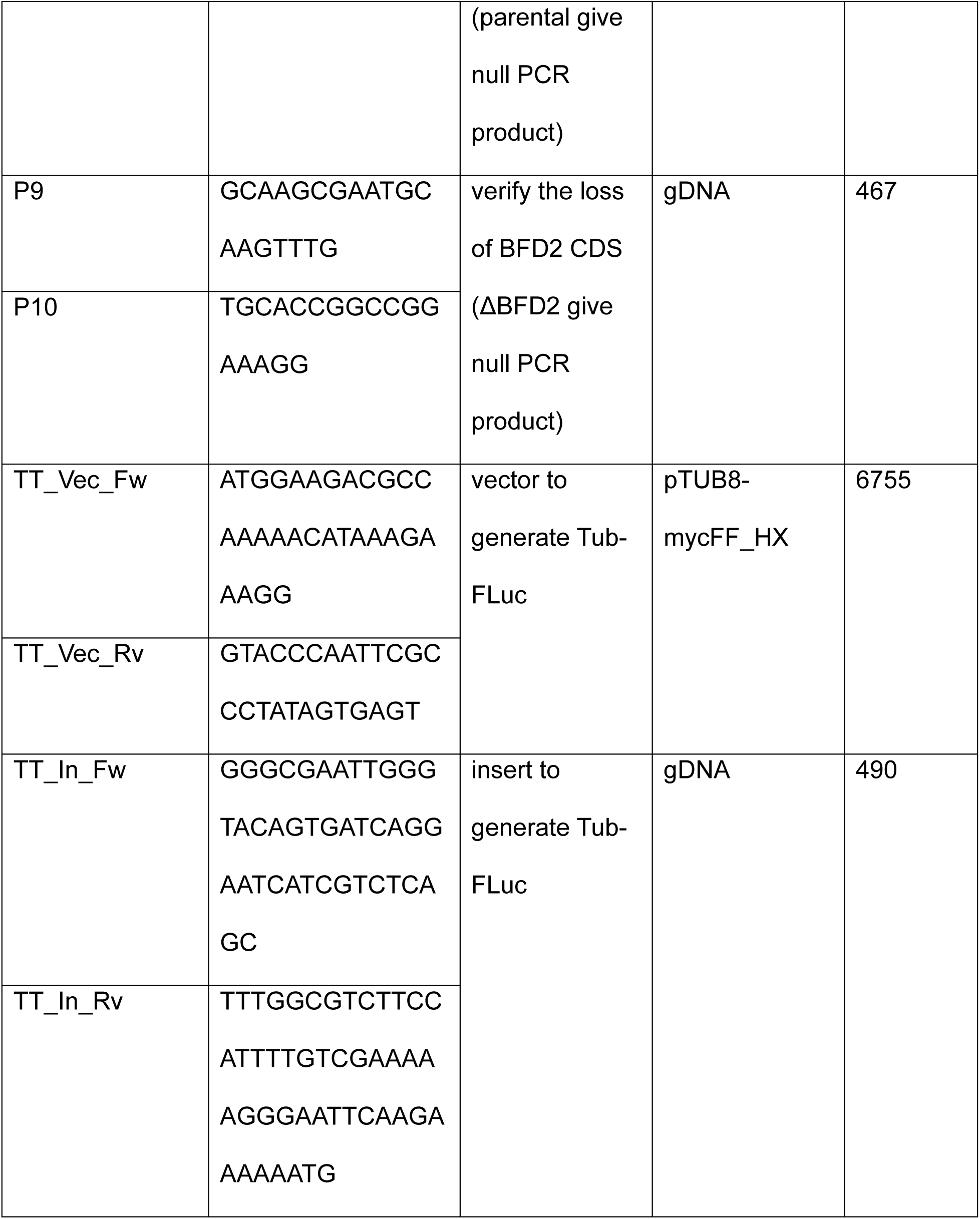
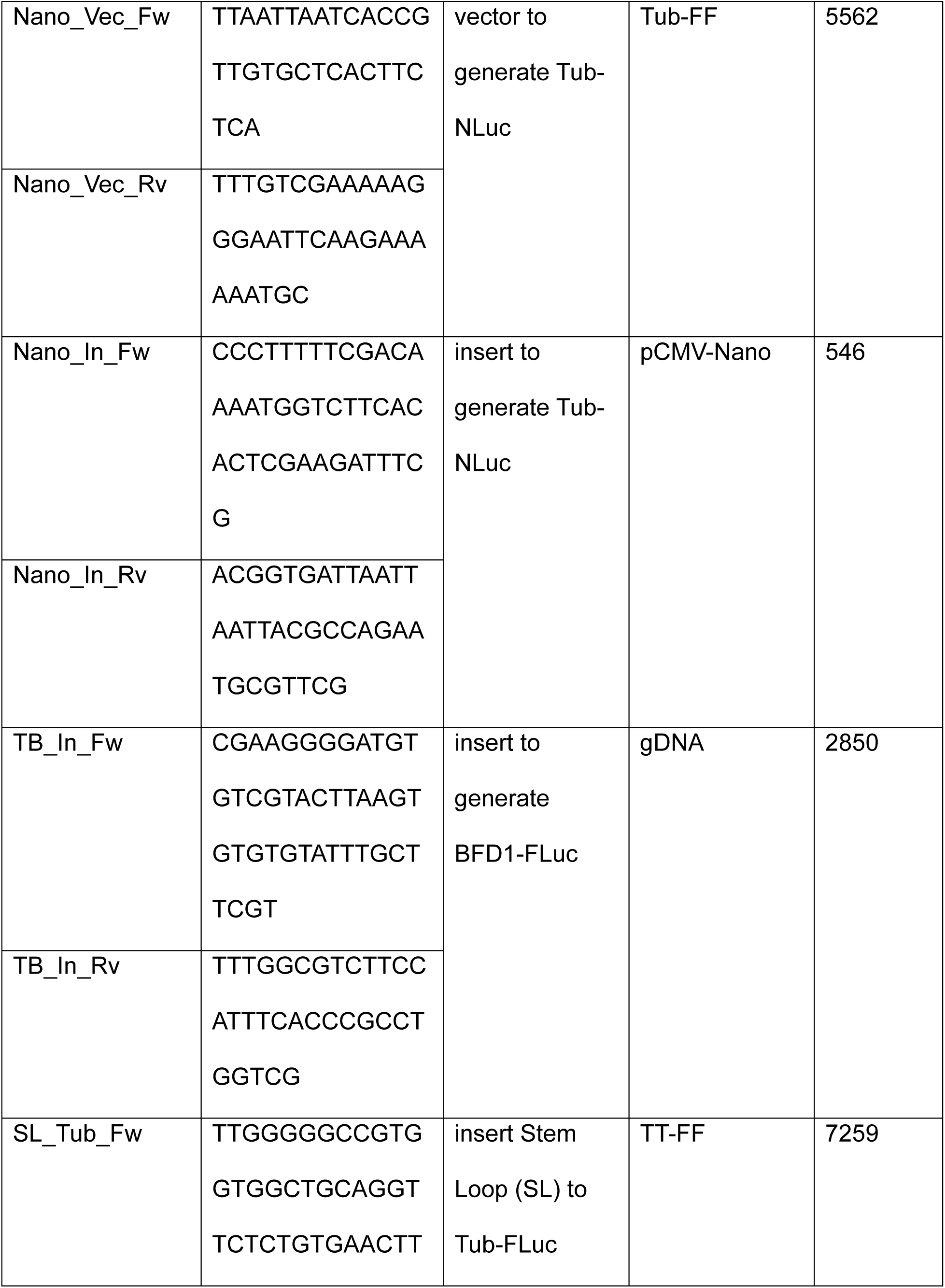
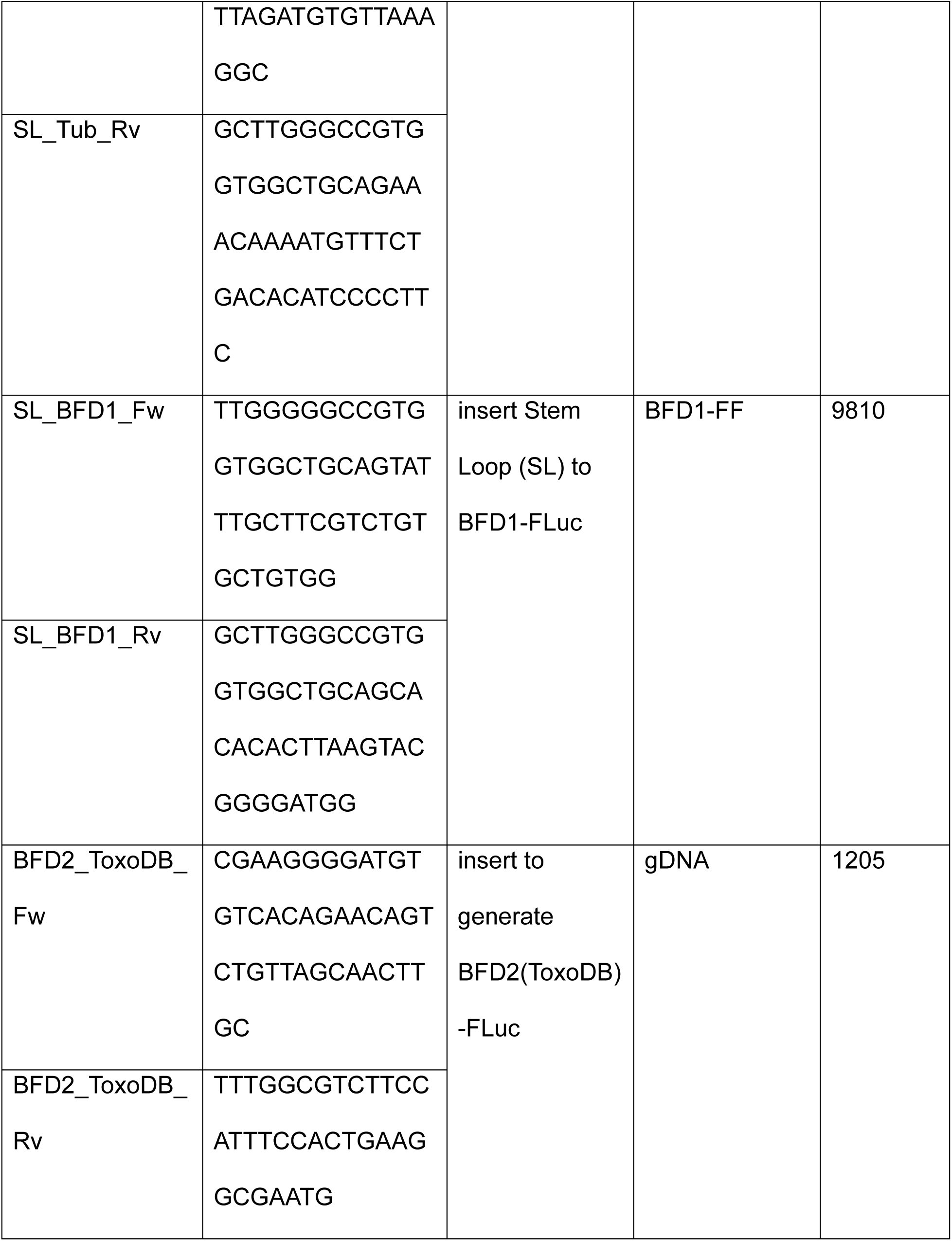
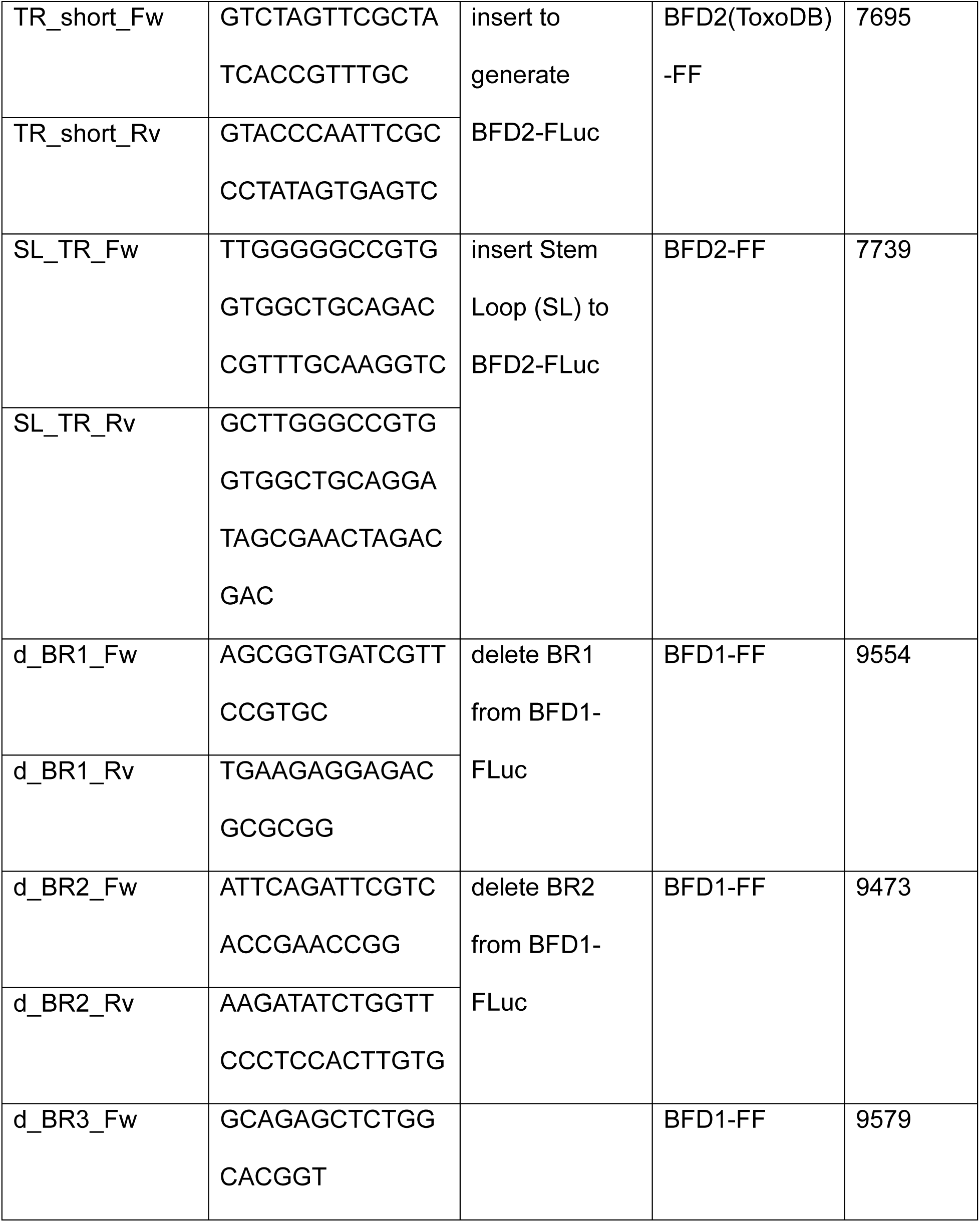
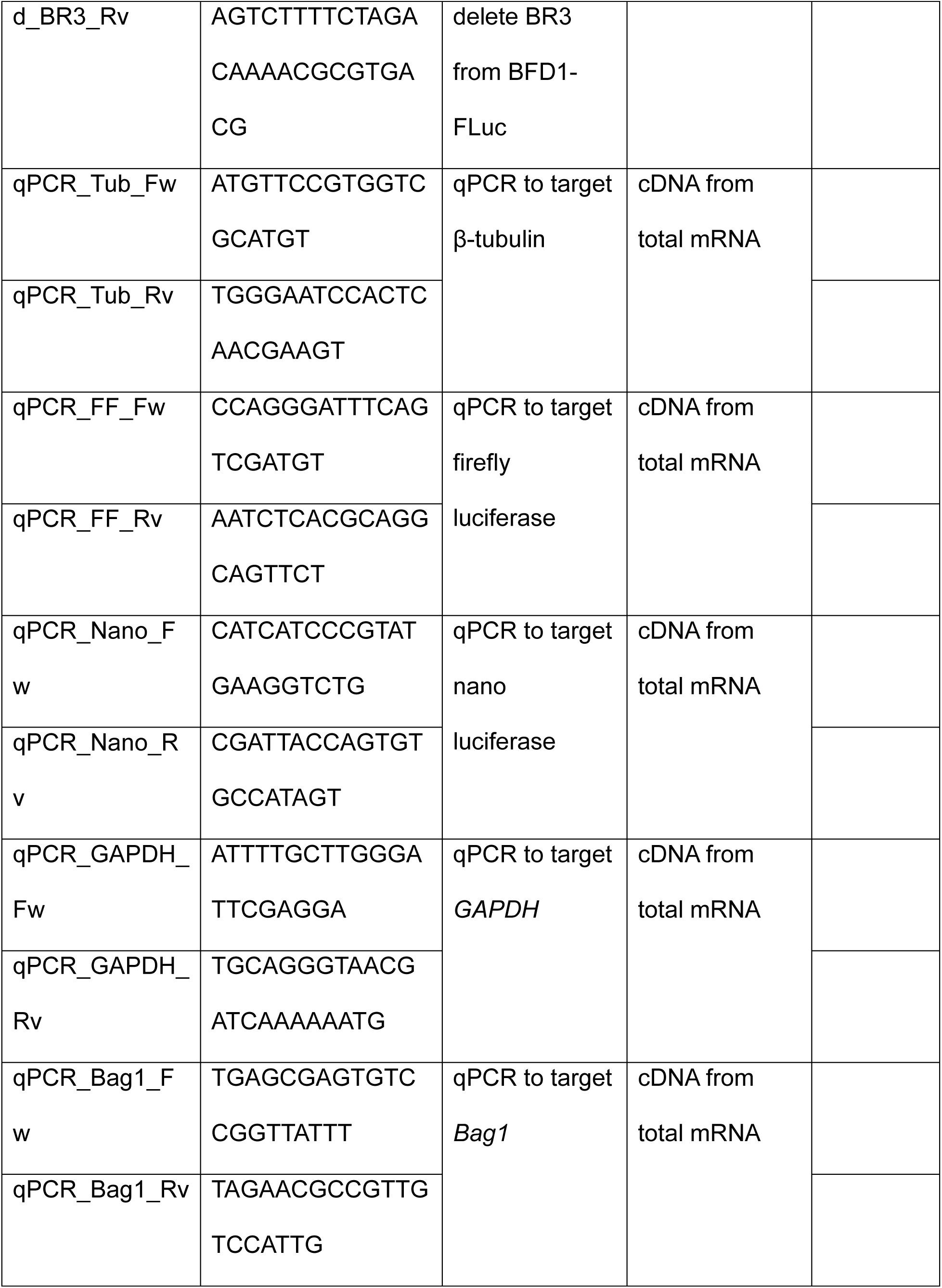
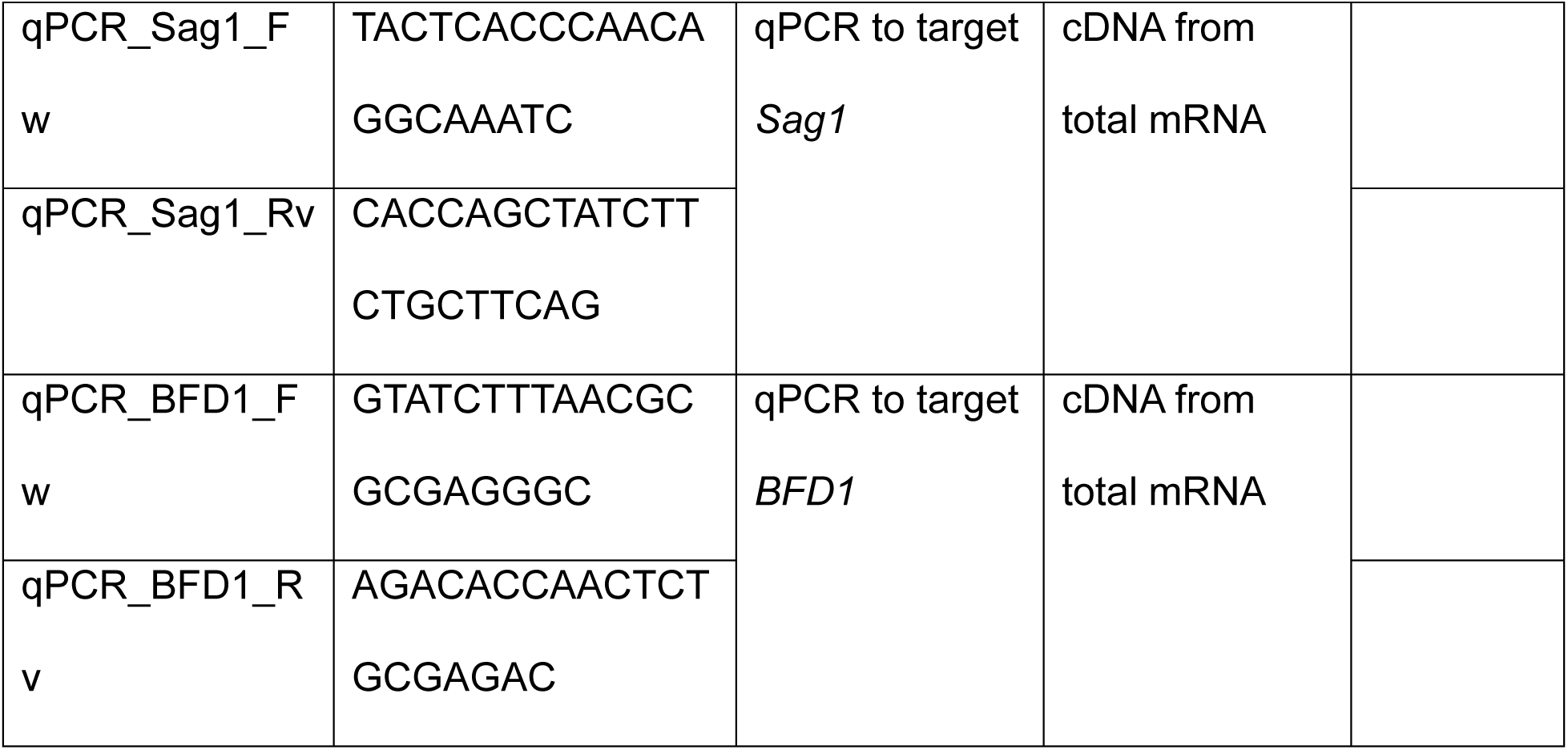
List of oligonucleotides used in this study.

