## Supplemental Figures for "Cap-independent translation directs stress-induced differentiation of the protozoan parasite *Toxoplasma gondii*"

William J. Sullivan Jr.^1,3*^

^1^Department of Pharmacology & Toxicology, ^2^Department of Biochemistry & Molecular Biology, ^3^Department of Microbiology & Immunology, Indiana University School of Medicine, Indianapolis IN

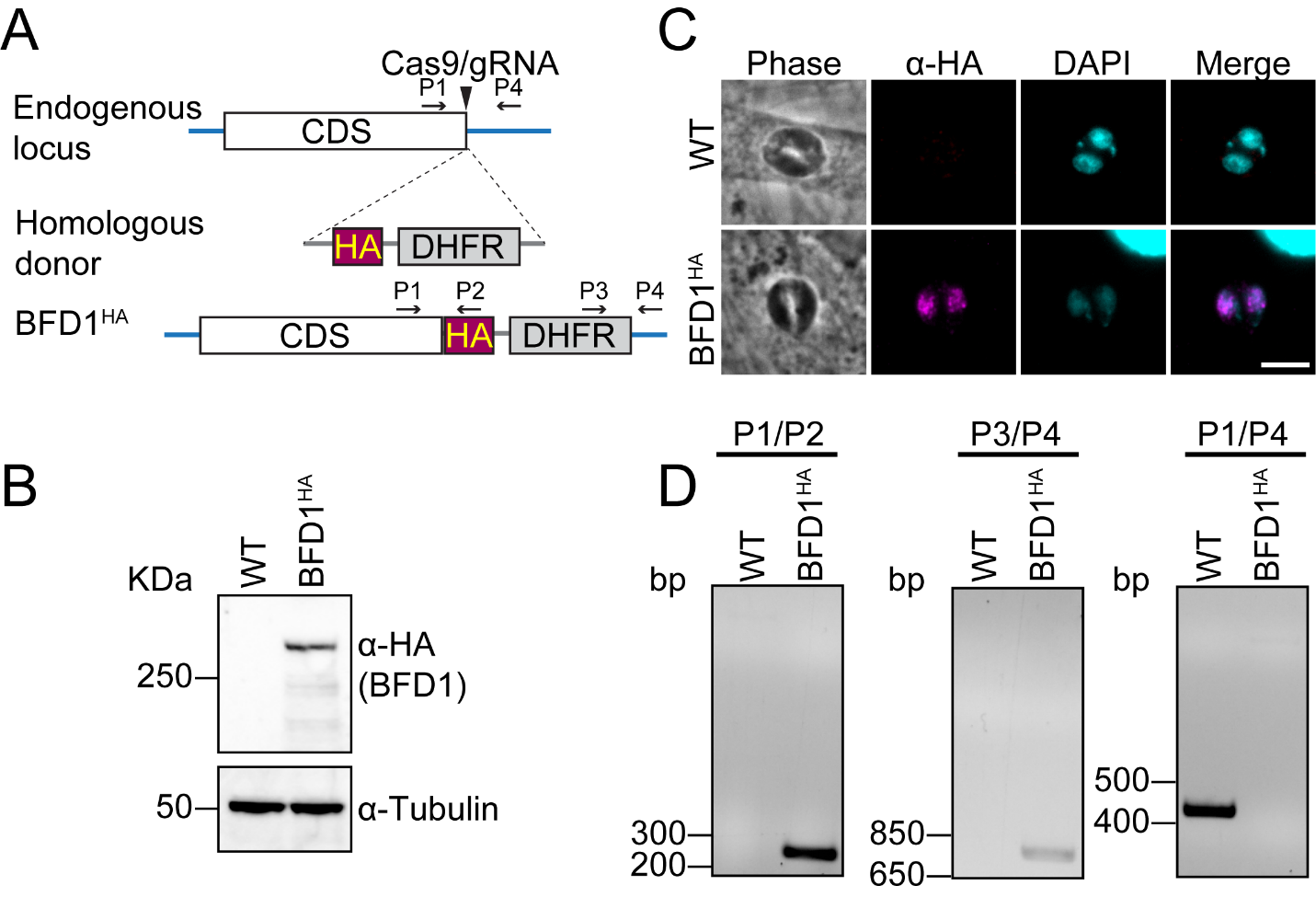

**Supplementary Figure 1. Endogenous tagging of BFD1^HA^.** (A) Schematic of the CRISPR/Cas9 approach to fuse endogenous BFD1 with a C-terminal HA epitope tag in ME49 strain parasites. DHFR served as a selectable marker. (B) Immunoblot of WT (parental) ME49 or BFD1^HA^ lysates after 24 h alkaline stress probed with anti-HA. Anti-tubulin was probed as a loading control. (C) IFA of WT ME49 and BFD1^HA^ parasites after 24 h alkaline stress using anti-HA (magenta). DAPI (blue) was used to visualize nuclei. Scale bar = 5 μm. (D) Diagnostic PCRs of gDNA were used to validate the proper integration of the HA tag using primers shown in (A).

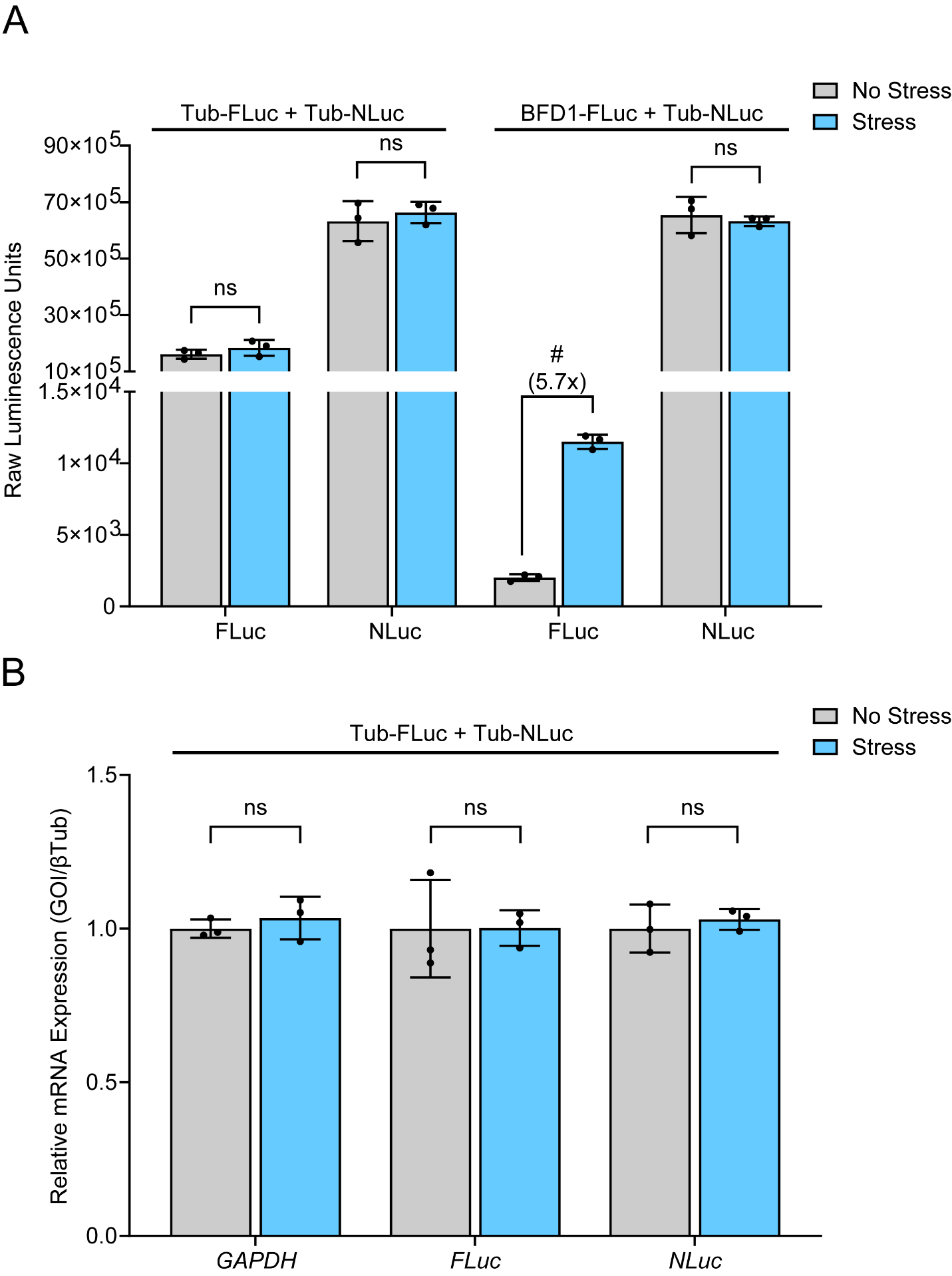

**Supplementary Figure 2. Alkaline stress does not affect NLuc activity or its mRNA levels.** The raw values of FLuc and NLuc measurements used to generate Fig. 1C are presented. (A) Luciferase activity in alkaline stress (Stress, blue bars) or tachyzoite conditions (No Stress, grey bars). Mean luciferase activity ± standard deviation from 3 biological replicates is plotted. ns = p>0.05; # = p≤0.0001 by Student’s two-tailed t-test. (B), Mean transcript abundance ± standard deviation from 3 biological replicates plotted relative to β-Tubulin with normalization to tachyzoites. GAPDH is included as an additional normalization to further support that β-Tubulin levels do not change in response to stress. ns = p>0.05 by Student’s two-tailed t-test. Fold changes are shown in parentheses.

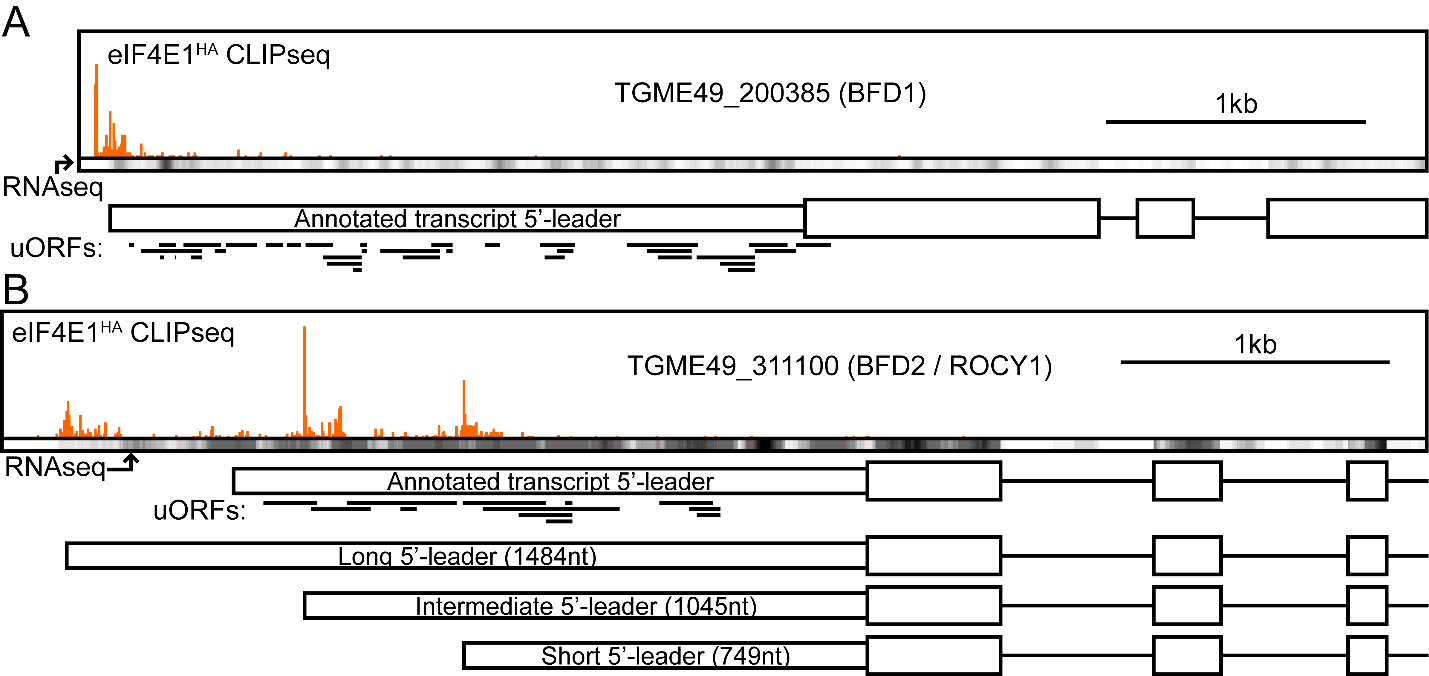

**Supplementary Figure 3. Transcriptional start site predictions of *BFD1* and *BFD2* transcripts.** eIF4E1-mRNA interaction sites as evidenced by CLIPseq are displayed as orange profiles above the gene models for *BFD1* (A) and *BFD2* (B). CLIPseq data was obtained from GSE243203. Strand-specific RNAseq heatmaps, obtained from GSE243206, are displayed below CLIPseq data to support transcript abundance over the gene bodies. 5’-leaders are displayed as narrow boxes, CDS as thick boxes, introns as lines. Predicted uORFs are displayed as thick black lines below the gene model. (A) The data presented is consistent with a single transcriptional start site corresponding to the annotated gene model for *BFD1*. (B) The data presented is inconsistent with the annotated *BFD2* 5’-leader and instead supports three transcript isoforms generated by alternate transcriptional start site usage, with a major 5'-leader ~750 nt in length used in the BFD2-FLuc reporter assays.

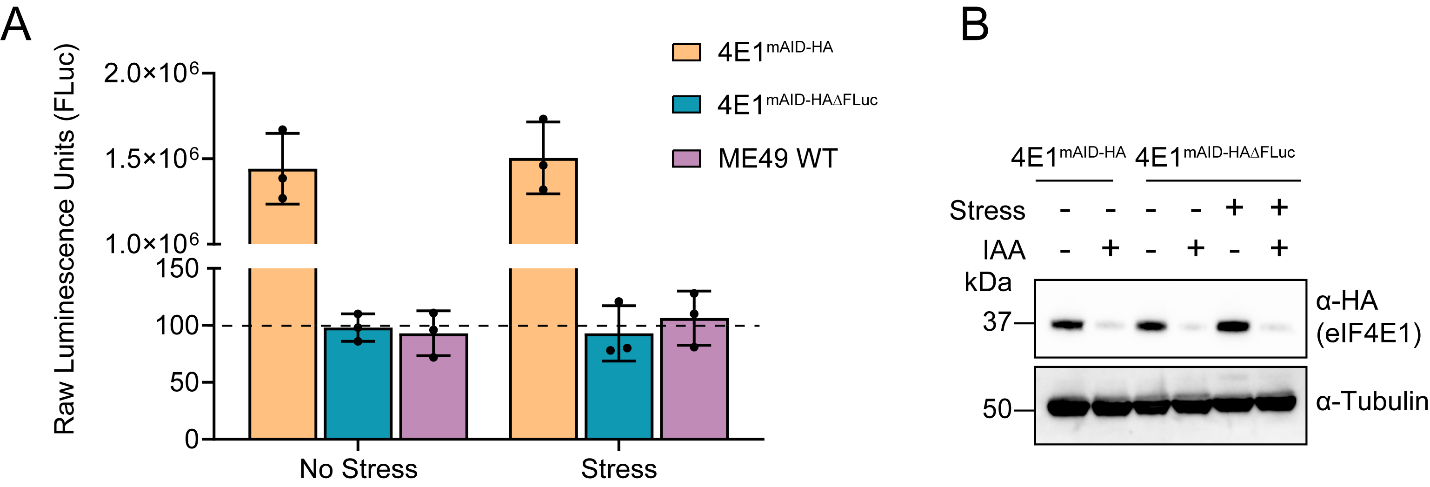

**Supplementary Figure 4. Knockout of FLuc in eIF4E1^mAID-HA^ parasites.** (A) Raw FLuc activity was measured for 10^6^ eIF4E1^mAID-HA^ parasites containing integrated FLuc and eIF4E1^mAID-HA^**^Δ^**^FLuc^ parasites in which the integrated FLuc gene was knocked out, under stress or unstressed conditions. The dotted line indicates the limit of luciferase activity detection in WT ME49 parasites as determined by taking the average from three independent measurements of untransfected parasites. (B) Immunoblot analyses with anti-HA to measure depletion of eIF4E1^mAID-HA^ in response to 4 h treatment with IAA in stress or unstressed conditions. Tubulin was probed as a loading control. Molecular weight markers are shown in kDa.

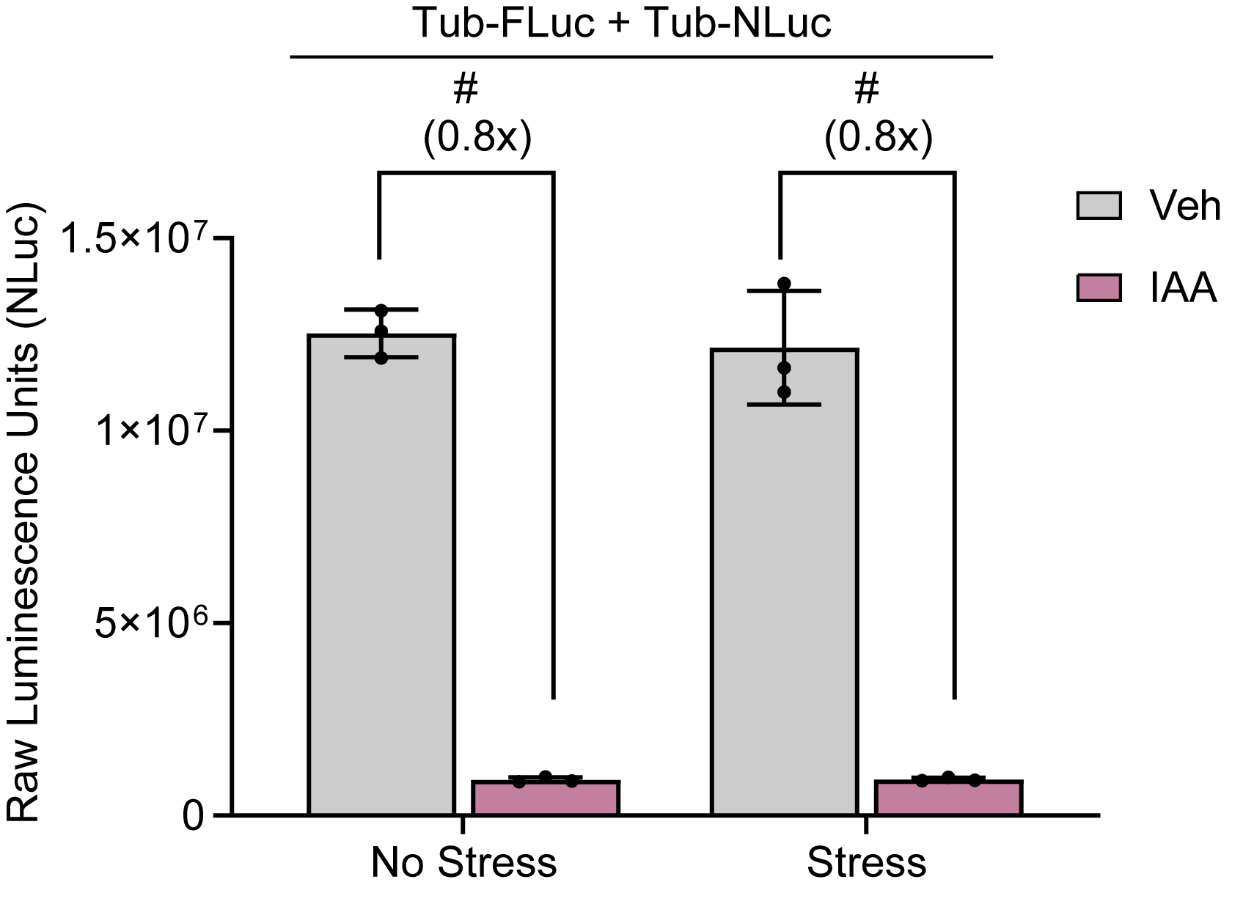

**Supplementary Figure 5. NLuc activity decreases in response to eIF4E1 depletion.** eIF4E1^mAID-HAΔFLuc^ parasites were co-transfected with Tub-FLuc and Tub-NLuc. Raw values of NLuc activity are shown. Bar graph represents mean NLuc activity ± standard deviation from 3 biological replicates. # = p≤0.0001 by Student’s two-tailed t-test. Fold changes are shown in parentheses.

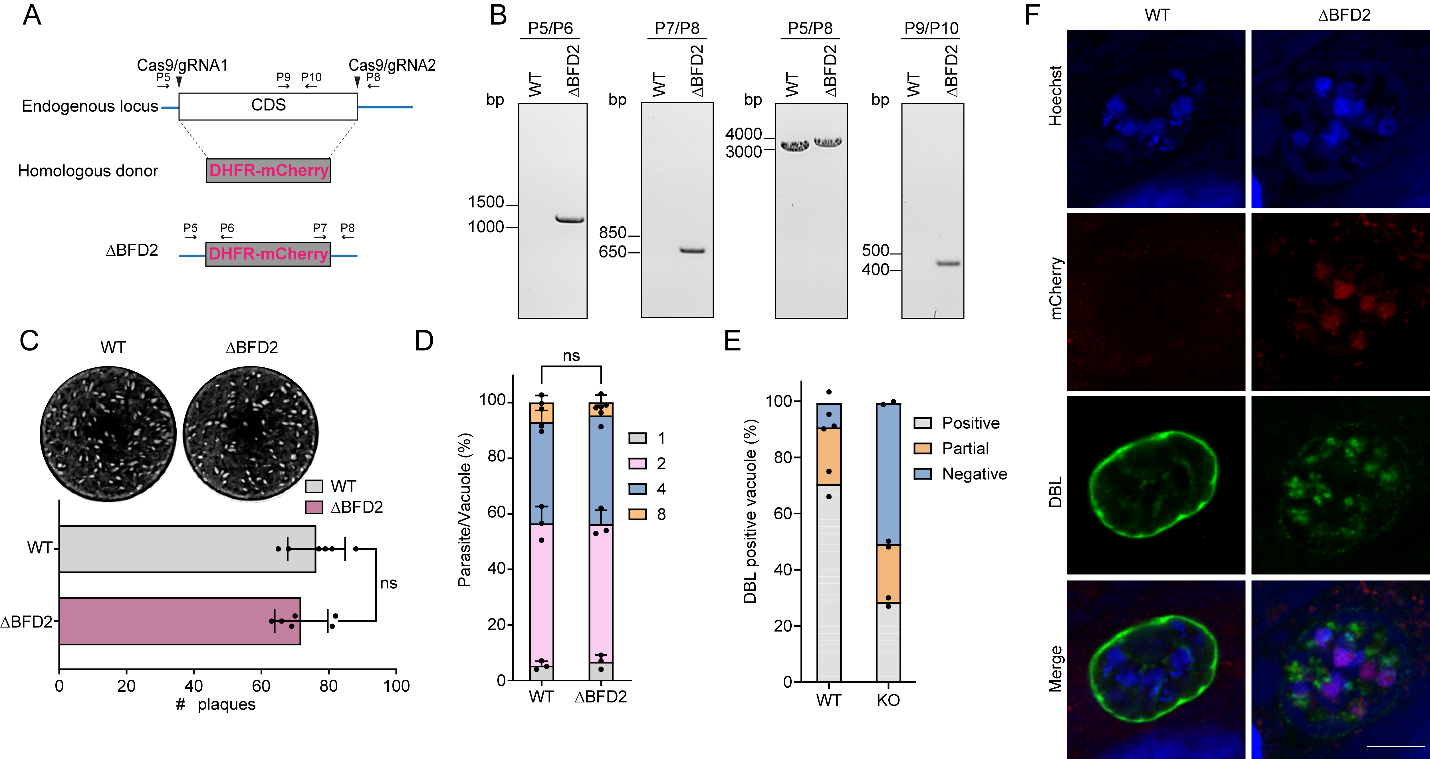

**Supplementary Figure 6. Generation and phenotyping of BFD2 knockout parasites.** (A) Schematic of CRISPR/Cas9 approach to generate ∆BFD2 parasites. DHFR-mCherry fusion operates as a selectable marker. (B) Genomic PCRs using primers shown in (A) that confirm replacement of the *BFD2* CDS with the DHFR-mCherry cassette. (C) Representative plaque assay of WT ME49 or ∆BFD2 parasites grown under tachyzoite culture conditions for 14 days. Quantitation of plaque number is included. ns = p>0.05 by Student’s two-tailed t-test. (D) Replication assay showing number of parasites per vacuole in WT ME49 and ∆BFD2 parasites 16 h post-invasion. Mean number of parasites per vacuole ± standard deviation is plotted from 3 biological replicates, with a minimum of 100 random vacuoles counted per sample. (E) Quantitation of parasite differentiation of WT ME49 and ∆BFD2 parasites after 72 h of alkaline stress from two biological replicates. Cultures were stained with FITC-labeled *Dolichos biflorus* lectin (DBL) to visualize cyst walls. A minimum of 100 random vacuoles were counted per sample. (F) Representative vacuoles counted in (E) after 72 h alkaline stress. DBL (green) stains cyst wall. mCherry (red) exclusively stains ∆BFD2 parasites. Hoechst (blue) used to visualize nuclei. Scale bar = 10 μm.

**Supplemental Table S1. List of oligonucleotides used in this study.**
